# Top-down attention and Alzheimer’s pathology impact cortical selectivity during learning, influencing episodic memory in older adults

**DOI:** 10.1101/2024.12.04.626911

**Authors:** Jintao Sheng, Alexandra N. Trelle, America Romero, Jennifer Park, Tammy T. Tran, Sharon J. Sha, Katrin I. Andreasson, Edward N. Wilson, Elizabeth C. Mormino, Anthony D. Wagner

**Affiliations:** Department of Psychology, Stanford University, Stanford, CA, USA; Department of Neurology & Neurological Sciences, Stanford University School of Medicine, Stanford, CA, USA; Wu Tsai Neurosciences Institute, Stanford University, Stanford, CA, USA

**Keywords:** episodic memory, neural selectivity, attention, Alzheimer’s disease, aging, functional MRI

## Abstract

Human aging affects the ability to remember new experiences, in part, because of altered neural function during memory formation. One potential contributor to age-related memory decline is diminished neural selectivity –– i.e., a decline in the differential response of cortical regions to preferred vs. non-preferred stimuli during event perception –– yet the factors driving variability in neural selectivity with age remain unclear. We examined the impact of top-down attention and preclinical Alzheimer’s disease (AD) pathology on neural selectivity during memory encoding in 156 cognitively unimpaired older participants who underwent fMRI while performing a word-face and word-scene associative memory task. Neural selectivity in face- and place-selective cortical regions was greater during events that were later remembered compared to forgotten. Critically, neural selectivity during learning positively scaled with memory-related variability in top-down attention, whereas selectivity negatively related to early AD pathology, evidenced by elevated plasma pTau_181_. Path analysis revealed that neural selectivity at encoding mediated the effects of age, top-down attention, and pTau_181_ on associative memory. Collectively, these data reveal multiple pathways that contribute to memory differences among older adults –– AD-independent reductions in top-down attention and AD-related pathology alter the precision of cortical representations of events during experience, with consequences for remembering.

## Introduction

The ability to remember experiences (i.e., episodic memory) is central to effective living, as memories inform understanding of ongoing events, shape thought, and guide decisions and action (*1*). Episodic memory declines later in the lifespan, concurrent with alterations in brain structure and function (*2*–*5*) related to (*6*–*8*) and independent of (*9, 10*) early Alzheimer’s disease (AD) pathology. At a representational level, the ability to subsequently remember an experience depends, in part, on establishing neural representations of the event’s features (or content) as the event unfolds, and extant data indicate that these cortical representations are altered in aging. In particular, neural selectivity –– the ability of neurons or brain regions to respond preferentially to specific stimuli or cognitive processes –– is lower in older than younger adults (i.e., neural dedifferentiation) and this lower selectivity is thought to contribute to age-related cognitive decline (*11*–*16*). While a well-established phenomenon, the factors that impact age-related and/or age-independent declines in neural selectivity remain unclear. Here, we use functional MRI (fMRI) along with biomarkers of preclinical AD pathology in a large sample of cognitively unimpaired (CU) older adults to identify mechanisms underlying neural selectivity during episodic memory formation in human aging.

In humans and other primates, top-down attention is known to influence neural selectivity during perception (*17*–*22*) and memory encoding (*23*–*26*), and predicts subsequent memory performance (*25, 27*–*30*). For example, top-down attention reduces the overlap in the neural populations representing attended vs. unattended objects, increasing neural selectivity in ventral visual cortex (*17*). Extant data further indicate that top-down attention is diminished in older relative to younger adults (*31*–*33*), and that age-related differences in attention partially account for age-related declines in episodic memory (*34, 35*). Yet, there is marked variability in attention-dependent performance (*36*) and in episodic memory (*8, 37*) across older adults. Here, we hypothesize that individual differences in the strength of neural selectivity across CU older adults may partially reflect differences in top-down attention, with consequences for subsequent memory.

Hallmark pathological features of AD – tau and amyloid-β (Aβ) proteins – begin accumulating decades before the emergence of clinical symptoms of dementia (a stage referred to as preclinical AD)(*38, 39*). Amyloid deposition is typically widespread throughout many neocortical association areas, including in frontoparietal cortical areas (*40*), whereas abnormal tau accumulation begins in the medial temporal lobe (*41, 42*), ventrolateral temporal cortex, and retrosplenial/posterior cingulate cortex (*43*–*45*). Importantly, this initial pattern of tau deposition is observed during the preclinical AD stage and is associated with reductions in memory performance (*42, 43, 46, 47*), raising the possibility that preclinical AD-related memory decline in aging may be partially mediated by changes in neural selectivity. Along these lines, Maass et al showed early tau PET burden was negatively associated with neural selectivity in the posterior-medial network (i.e., place-selective regions) in 50 CU older adults (*48*). Here, we leverage biofluid biomarkers of preclinical AD to explore the relations between AD pathology, neural selectivity, and memory in a large sample of CU older adults.

Moreover, to the extent that there are links between differences in attention and neural selectivity in older adults, an additional open question is whether such links relate to preclinical AD processes. On the one hand, extant data indicate that the locus coeruleus, one of the earliest brain regions affected by tau protein accumulation (*49, 50*), plays a crucial role in arousal and alerting (*51*), and modulates multiple aspects of attentional function through interactions with the dorsal attention network (DAN) (*52, 53*). Consequently, top-down up attention, supported by DAN (*54, 55*), may be influenced by early AD pathology (*56, 57*). On the other hand, age-related changes in neural systems of attention may occur through AD-independent processes, including processes that give rise to structural (*58*–*60*) and functional (*61, 62*) changes to the DAN. As such, we examine whether early AD pathology impacts neural selectivity in CU older adults through disease-related changes in attention, with consequences for later remembering, or whether any links between AD pathology, neural selectivity, and memory represent a distinct pathway from that related to attention.

To address these knowledge gaps, we leverage a large sample (n = 156) of CU older adults (60 – 88 years) from the Stanford Aging and Memory Study (SAMS) who performed a word-face and word-place associative memory task (**Fig. 1A**) concurrent with high-resolution fMRI (*8, 37*). Here, we focus on neural selectivity during memory formation, defined as univariate differences in BOLD signal between preferred vs. non-preferred stimuli in face- and place-selective regions-of-interest (ROIs). Additionally, we define univariate activation differences between subsequently remembered vs. forgotten trials (i.e., subsequent memory effects; SME) in the DAN as a memory-relevant neural index of top-down attention (*54, 55, 63*) and, in a subset of participants, we assay early AD pathology using plasma (n = 138) and CSF (n = 115) pTau_181_ and Aβ_42_/Aβ_40_. Analyses examine whether trial-level and subject-level measures of neural selectivity during encoding predict subsequent memory; how neural selectivity varies with age, top-down attention, and AD biomarkers; and, using structural equation modeling (SEM), whether multiple pathways account for memory variability among CU older adults.

**Fig. 1.**
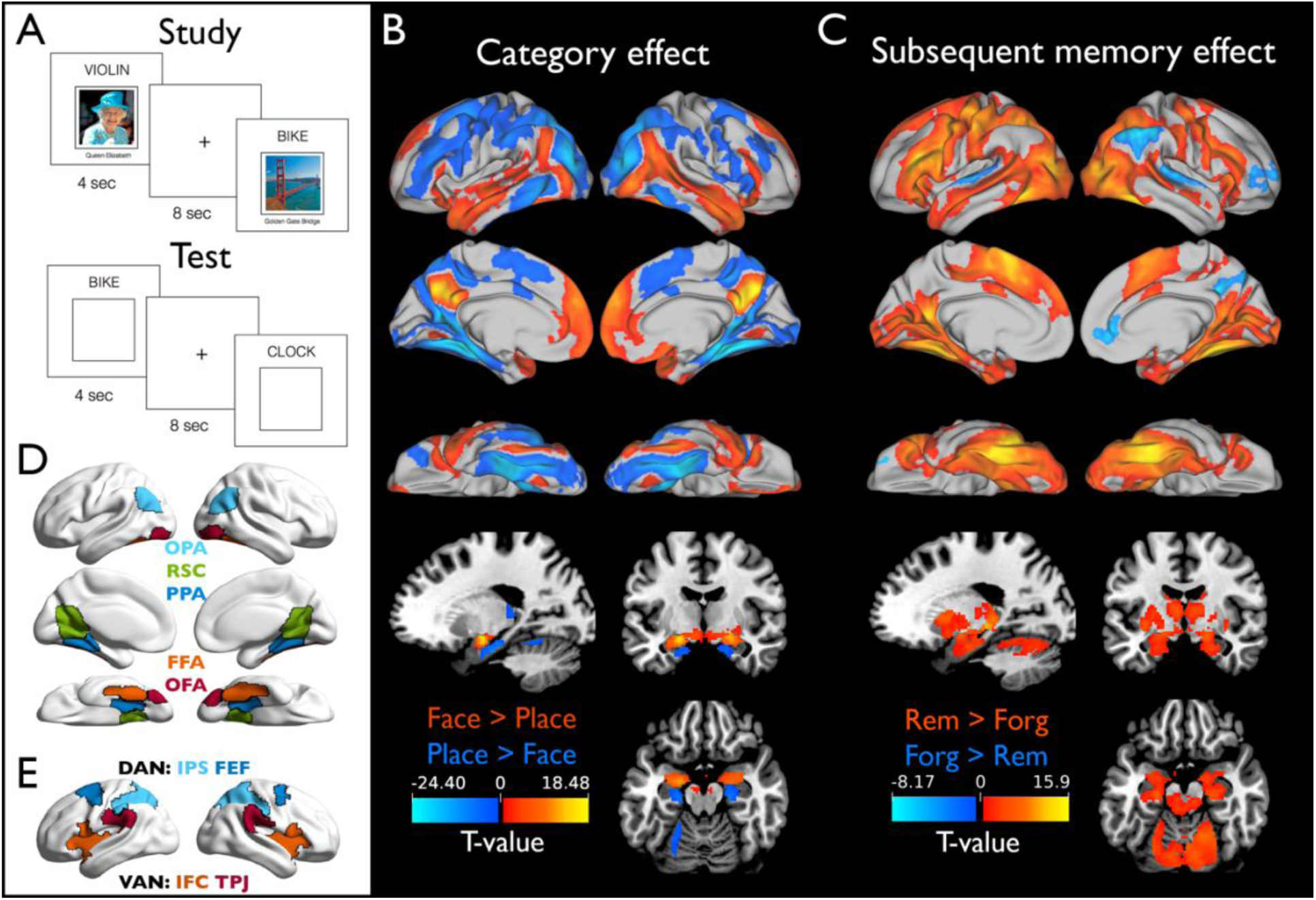
Experimental design, group-level statistical maps, and regions of interest (ROIs). (A) Structure of study (encoding) and test (retrieval) trials in the word-picture associative memory task. Group-level category effects (face vs. place) during word-picture study. (C) Group-level subsequent memory effects depicting differential activation on word-picture encoding trials for which the association was subsequently remembered vs. subsequently forgotten at test. (D) Predefined place- and face-selective ROIs: OPA, occipital place area; RSC, restrosplenial cortex; PPA, parahippocampal place area; FFA, fusiform face area; OFA, occipital face area. (E) Predefined frontoparietal dorsal attention network (DAN) and ventral attention network (VAN) ROIs: IPS, intraparietal sulcus; FEF, frontal eye fields; IFC, inferior frontal cortex; TPJ, temporoparietal junction.

## Results

### Associative memory behavior

Associative *d’* (*Z*_associative hit rate_ − *Z*_associative false alarm rate_) assessed memory performance, computed overall (i.e., pooled across categories) and separately for face (i.e., word-face associations) and place (i.e., word-place associations) memory, declined with age (**Fig. S1**; overall: β = -.044, *p* < .001; face: β = -.030, *p* = .003; place: β = -.044, *p* < .001). A marginal age × category interaction suggests that age may impact memory for places more than for faces (*F*_1, 164_ = 3.477, *p* = .064).

### Category and subsequent memory effects

Contrasting word-face vs. word-place encoding trials revealed well-established neural category effects (**Fig. 1B; Table S1**). Specifically, activation was higher on face trials in bilateral occipital face area (OFA), fusiform face area (FFA), precuneus, superior temporal sulcus, anterior temporal lobe, medial prefrontal cortex, and amygdala, amongst other regions, whereas activation was higher on place trials in bilateral occipital place area (OPA), parahippocampal place area (PPA), retrosplenial cortex (RSC), dorsolateral prefrontal cortex (dlPFC), and anterior hippocampus, amongst other regions.

To identify associative encoding effects, we conducted a subsequent memory analysis(*64, 65*) that contrasted remember (Rem) trials, consisting of word-picture trials that were later remembered (i.e., subsequent associative hits), with forgotten (Forg) trials, consisting of all other study trials (i.e., subsequent associative misses/incorrect category response, subsequent item hits/‘old’ response, and subsequent item misses/‘new’ response). Activation was higher during subsequently remembered than forgotten trials in bilateral ventral visual cortex, frontoparietal components of the DAN, ventral attention (VAN), and default mode (DMN) networks, and hippocampus, amongst other regions (**Fig. 1C; Table S2**). Greater activation on subsequently forgotten vs. remembered trials was observed in right angular gyrus, precuneus, anterior cingulate cortex/medial prefrontal cortex, dlPFC/frontopolar cortex, and bilateral superior temporal gyrus.

### Subsequently remembered trials show greater neural selectivity at encoding

For ROI-based analyses, we generated five predefined ROIs known to demonstrate face-selective (FFA and OFA) or place-selective activation (RSC, PPA, and OPA) (**Fig. 1D**) and using a leave-one-out procedure to ensure statistical independence (**Fig. S2**; *Methods*). We then extracted the mean univariate effect (i.e., mean *t*-value) in these ROIs when contrasting preferred vs. non-preferred categories (i.e., neural selectivity) as a function of subsequent memory.

Neural selectivity in place-(PPA, OPA, and RSC) and face-selective (FFA and OFA) ROIs was greater than zero (*t*s = 8.04 – 27.94, *p*s_Holm_ < .001; **Fig. S3A**), confirming each exhibits greater activity during encoding of words paired with images of the ROI’s preferred category. Moreover, neural selectivity at encoding was higher on subsequently remembered than forgotten trials in both place- and face-selective regions (*t*s = 6.62 – 11.02, *p*s_Holm_ < .001; **Fig. S3A**). As neural selectivity was strongly correlated across corresponding category-selective ROIs (**Fig. S3B**), we used mean selectivity across the three place-selective ROIs and across the two face-selective ROIs in all subsequent analyses.

### Age-related decreases in neural selectivity

A linear mixed-effect model (LMM) (**Fig. 2A, Table S3**), with age, memory (Rem vs. Forg), and region (Face-vs. Place-selective ROI) predictors, revealed that neural selectivity declined with age (*p* = .007), was greater on subsequently remembered trials (*p* < .001), and was greater in place-selective regions (*p* = .004). Moreover, age × region (*p* = .010) and age × memory (*p* = .001) interactions revealed that age-related decreases in neural selectivity were greater in place-(*β* = -.020, *p*_Holm_ < .001) compared to face-selective regions (*β* = -.004, *p*_Holm_ = .508) and on subsequently remembered (*β* = -.022, *p*_Holm_ < .001) compared to forgotten trials (*β* = -.002, *p*_Holm_ = .758). Finally, age-related declines in neural selectivity as a function of memory were similar across place- and face-selective ROIs (age × memory × region: *p* = .57). Collectively, these results demonstrate that neural selectivity is negatively associated with age, particularly for subsequently remembered trials and place-selective regions.

**Fig. 2.**
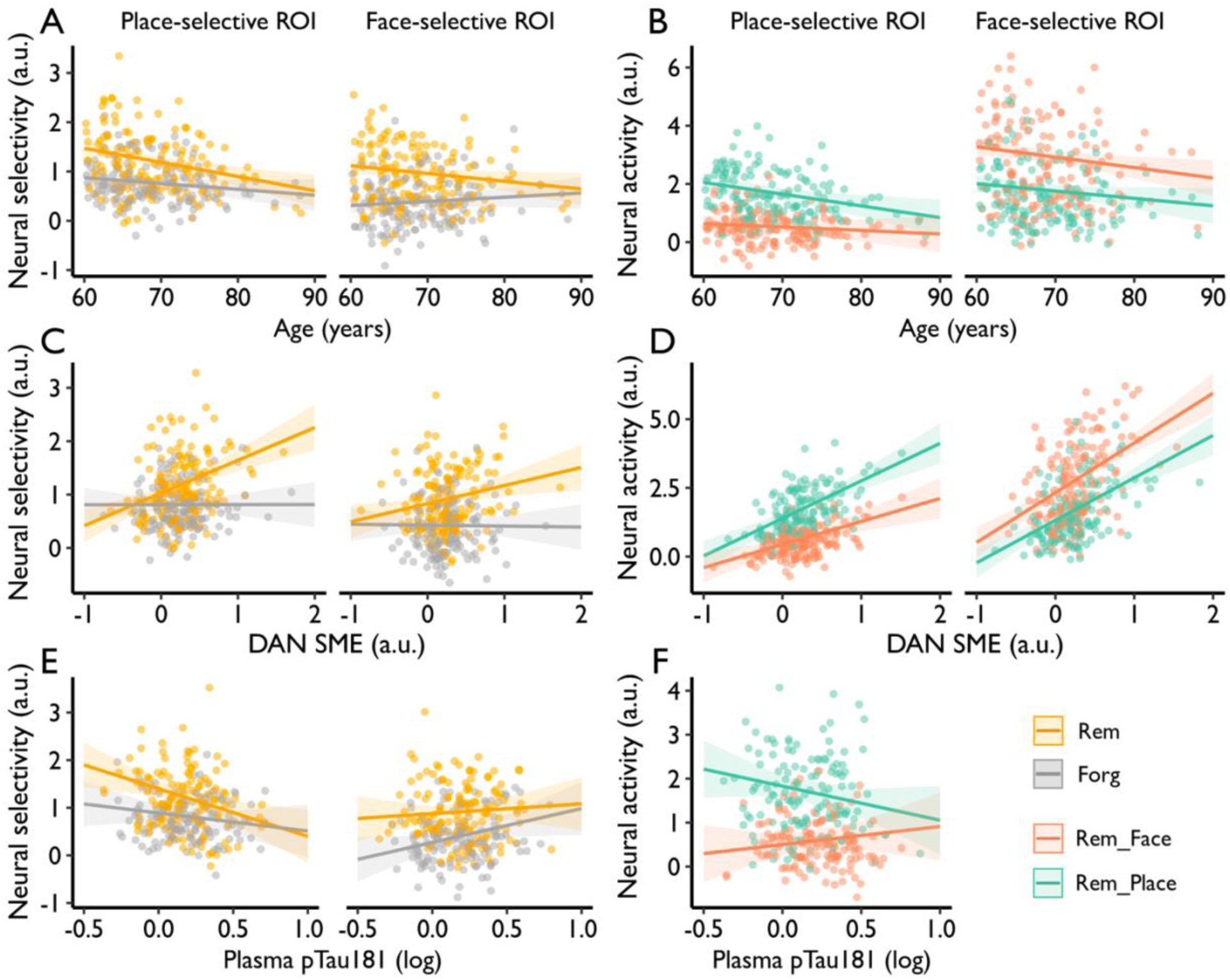
Neural selectivity relates to age, top-down attention, and early AD pathology. (A) The age-related decrease in neural selectivity at encoding is greater on subsequently remembered vs. forgotten trials. (B) Neural activity decreased more with age for preferred category than non-preferred category trials in (left) place-but not (right) face-selective regions (analyses restricted to subsequently remembered trials). Sex and years of education were included as nuisance variables. (C) Neural selectivity on remembered trials was associated with the subsequent memory effect (SME) in frontoparietal nodes of the dorsal attention network (DAN). (D) DAN SME was tightly related to face- and place-related activity on subsequently remembered trials in (left) place- and (right) face-selective regions. (E) Neural selectivity on remembered trials in (left) place-but not (right) face-selective regions significantly declined with plasma pTau_181_. (F) Plasma pTau_181_ negatively related to preferred neural activity (i.e., place-related activity, cool color) in place-selective regions. Age, sex, and years of education were included as nuisance variables.

To determine if age-related reductions in selectivity reflect attenuation (decreased activity for preferred stimuli) and/or broadening (increased activity for non-preferred stimuli) (*66*), we examined preferred- and non-preferred-category activity in face- and place-selective ROIs, restricting analyses to remembered trials (n.b., selectivity did not change on forgotten trials). An age × category (face vs. place) × region (Face-vs. Place-selective ROI) interaction (*p* = .028; **Table S4**) indicated that the pattern of age-related change in selectivity differed across face- and place-selective ROIs. Follow-up analyses revealed an age × category interaction in place-selective regions (*p* = .023), indicating that the age-related decline in neural selectivity was driven by decreased place/preferred-category activity (*β* = -.040, *p*_Holm_ = .013), without a corresponding increase in face/non-preferred-category activity (*β* = -.012, *p*_Holm_ = .381) (**Fig. 2B**, left). In face-selective regions, the interaction was not significant (*p* = .405); face/preferred-category activity decreased with age (*β* = -.036, *p*_Holm_ = .028) and a qualitatively similar pattern was seen for place/non-preferred-category activity (*β* = -.025, *p*_Holm_ = .128) (**Fig. 2B**, right). As such, age-related declines in neural selectivity reflect attenuation rather than broadening, with differential reductions in preferred vs. non-preferred activity being more evident in place-selective regions (potentially because face-selective regions show above-baseline activity for both categories).

### Neural selectivity is modulated by top-down attention

Attenuation of neural selectivity with age and its link to subsequent memory may reflect reduced top-down attentional modulation of category-selectivity activity. To explore this possibility, we extracted the mean univariate effect (i.e., mean *t*-value) when contrasting subsequently remembered and forgotten trials (i.e., SME) in frontoparietal regions of the DAN (**Fig. 1E**; see *Methods*). We observed a significant SME in DAN (*t*_155_ = 9.43, *p* < .001) that did not significantly vary with age (**Fig. S4A**; *β* = -.006, *p* = .176).

An LMM (**Fig. 2C, Table S5**), with top-down attention (DAN SME), memory (Rem vs. Forg), and region (Face-vs. Place-selective ROI) as predictors of neural selectivity, revealed an effect of top-down attention (*p* = .002) and a top-down attention × memory (*p* < .001) interaction. These results reflect that neural selectivity increased with greater memory-related top-down attention (DAN SME), and this association was evident for later remembered (*β* = .476, *p*_Holm_ < .001) but not forgotten trials (*β* = -.008, *p*_Holm_ = .929). Effects of memory-related top-down attention were similar in place- and face-selective regions (top-down attention × memory × region: *p* = .252; top-down attention × region: *p* = .189). Notably, when age was included in the model, we found that DAN SME (*p* = .002) and age (*p* = .016) explained unique variance in neural selectivity (**Table S5**). Parallel analysis with frontoparietal regions of the VAN, which is linked to bottom-up attention (*54, 55*) (**Fig. 1E**), revealed that these effects were specific to DAN-mediated top-down attention rather than reflecting a more general impact of attentional processes on neural selectivity (**Fig. S4B**; ***Supplementary Results***). Thus, during encoding trials that were later remembered, neural selectivity in content-sensitive cortex scaled with memory-related frontoparietal top-down attention.

To determine whether the association between DAN activity and neural selectivity reflects modulation of activity for the preferred and/or non-preferred stimulus category, we decomposed selectivity on remembered trials into face and place activity. An LMM revealed a top-down attention (DAN SME) × category (Face vs. Place) × region (Face-vs. Place-selective regions) interaction (*p* = .010; **Table S6**). Follow-up analyses revealed a DAN SME × category interaction in place-(*p* = .016) but not face-selective regions (*p* = .224), which reflects a stronger association between DAN SME and neural activity for place/preferred (*β* = 1.365, *p*_Holm_ < .001) than face/non-preferred trials (*β* = .838, *p*_Holm_ < .001) in place-selective regions, but similar associations for face/preferred (*β* = 1.811, *p*_Holm_ < .001) and place/non-preferred trials (*β* = 1.546, *p*_Holm_ < .001) in face-selective regions (**Fig. 2D**). Thus, on subsequently remembered trials, top-down attention increased sensitivity to both preferred and non-preferred stimuli, impacting neural selectivity in place-selective regions by differentially enhancing responses to preferred stimuli.

### Neural selectivity declines with early AD pathology

Early AD biomarkers of Aβ_42_/Aβ_40_ and pTau_181_, measured in blood plasma (n = 138) and CSF (n = 115), varied with age (***Supplementary Results***). Controlling for age, sex, and education, an LMM predicting neural selectivity (**Table S7**) revealed a plasma pTau_181_ × region (Face-vs. Place-selective ROI) interaction (*p* < .001), with higher plasma pTau_181_ related to decreased neural selectivity in place-(*β* = -.690, *p* = .011) but not in face-selective regions (*β* = .458, *p* = .090) (**Fig. 2E**). While the plasma pTau_181_ × memory (Rem vs. Forg) interaction in place-selective regions was not significant (*F*_1,405_ = 2.164, *p* = .142), neural selectivity on remembered trials declined with pTau_181_ (Rem: *β* = -1.0, *p* = .004; Forg: *β* = -.379, *p* = .268). A similar pattern of decreased neural selectivity in place-selective regions on remembered trials was found for plasma Aβ_42_/Aβ_40_ (**Fig. S5 and Table S7**). When decomposing neural selectivity in place-selective regions into activation on place and face trials, an LMM (**Table S8**) revealed a plasma pTau_181_ × category (Face-vs. Place) interaction (*p* = .002), with activation to place/preferred trials qualitatively declining (*β* = -.775, *p* = .102), but activation to face/non-preferred trials qualitatively increasing: *β* = .412, *p* = .384) (**Fig. 2F**). Thus, while selectivity in place-selective regions declined with plasma AD-related proteins, our findings are ambiguous with respect to whether this selectivity change with early AD pathology reflects attenuation and/or broadening.

Corresponding analyses of neural selectivity’s relationship with CSF pTau_181_ and Aβ_42_/Aβ_40_ (**Table S9**) revealed similar trends as those observed with plasma (**Fig. S6**), but the effects did not reach significance, perhaps due to the smaller sample size. Consistent with this possibility, restricting analyses to the participants with both plasma and CSF assays revealed qualitatively similar patterns to those reported above (compare **Fig. 2E** to **Fig. S7**). Finally, in contrast to neural selectivity, the DAN SME did not vary with plasma (pTau_181_: *β* = -.025, *p* = .918; Aβ_42_/Aβ_40_: *β* = 2.608, *p* = .229) nor CSF biomarkers (pTau_181_: *β* = .052, *p* = .778; Aβ_42_/Aβ_40_: *β* = -.786, *p* = .577), suggesting that memory-relevant variability in top-down attention is independent of early AD pathology in CU older adults.

### Top-down attention and AD biomarkers mediate age-related reductions in neural selectivity

Given observed associations between neural selectivity and age, plasma pTau_181_, and DAN activity, we conducted a mediation analysis to determine if plasma pTau_181_ and DAN SME mediated the effect of age on neural selectivity in place-selective cortex on remembered trials, with sex and years of education as confounding variables. Results revealed that both plasma pTau_181_ and DAN SME partially mediated the negative relationship between age and neural selectivity (Indirect1-pTau_181_: *a_1_b_1_* = -.007, *β*_standardized_ = -.066, 95% CI = [-.013, -.002]; Indirect2-DAN-SME: *a_2_b_2_* = -.005, *β*_standardized_ = -.046, 95% CI = [-.011, -.0003]; direct effect: *c’* = -.022, *β*_standardized_ = -.211, *p* = .002) (**Fig. 3A**). Moreover, comparison of the effect sizes in the two indirect pathways revealed that they did not differ from each other (Δ*β*_Indirect1-Indirect2_ = -.002, 95% CI = [-.010, .006]), suggesting that early AD pathology and top-down attention have comparable, independent effects on age-related declines in neural selectivity.

**Fig. 3.**
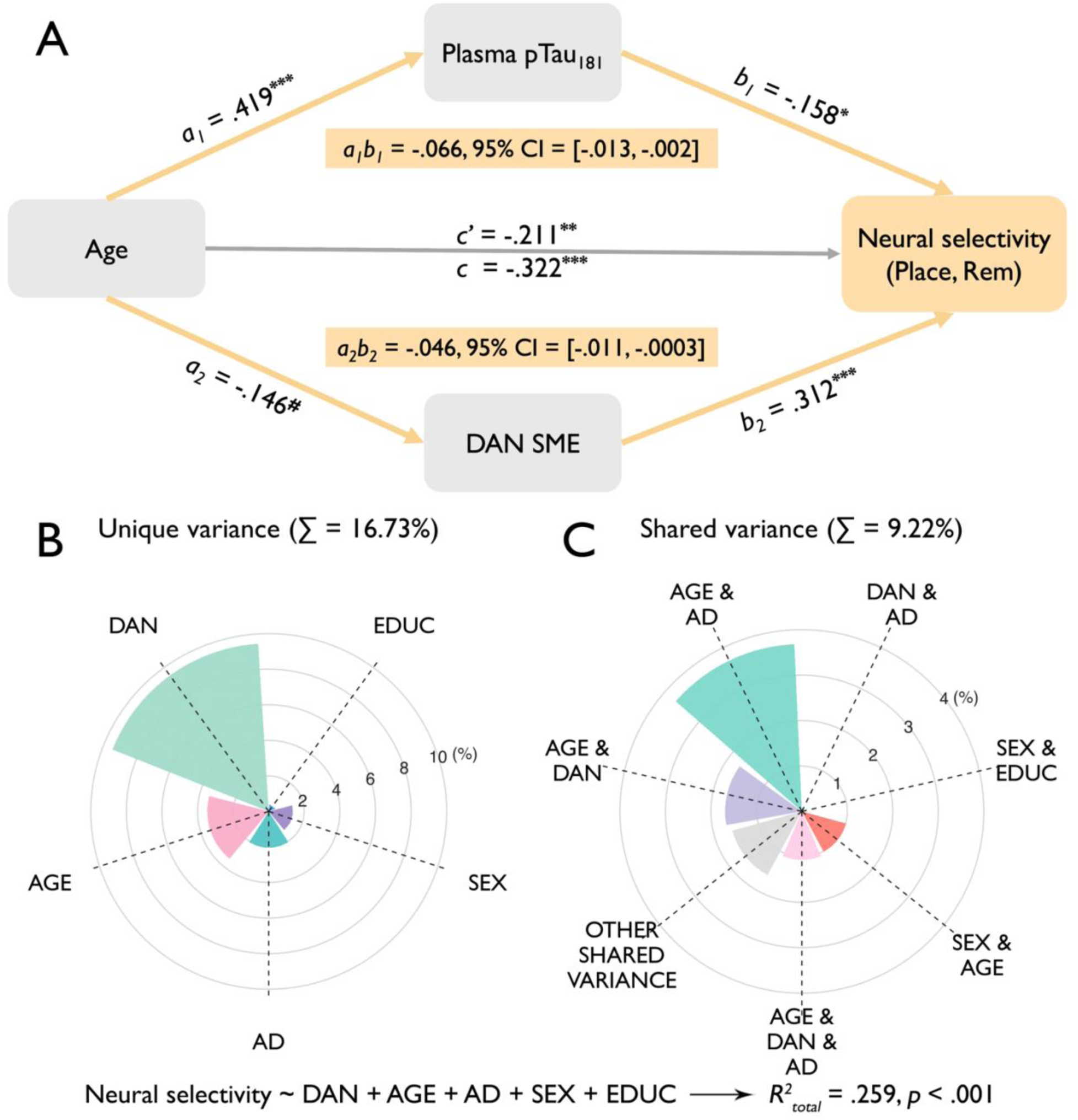
Predictors on neural selectivity. (A) Plasma pTau_181_ and DAN SME partially mediated the negative relationship between age and neural selectivity (on remembered trials in place-selective regions). Sex and years of education were included as nuisance variables. **^#^** *p* = .05, **\*** *p* < .05, **\*\*** *p* < .01, **\*\*\*** *p* < .001. *c’* = direct effect; *c* = total effect = *a_1_b_1_ + a_2_b_2_ + c’*. (B) Unique and (C) shared explained variances of predictors of neural selectivity. DAN = DAN SME (i.e., subsequent memory effect in dorsal attention network), AD = plasma pTau_181_, EDUC = years of education.

To further summarize the unique and shared contributions of age, DAN SME, and plasma pTau_181_ to neural selectivity, we combined these factors in a multiple linear regression model predicting selectivity in place-selective cortex on remembered trials, with sex and education included as covariates. This model explained 25.9% total variance in selectivity (*p* < .001; **Table S10**), with DAN SME explaining the most unique variance (*R^2^*_DAN_ = *R^2^* - *R^2^* = 9.4%), followed by age (*R^2^* = *R^2^*-*R^2^* = 3.4%), and plasma pTau_181_ (*R^2^*_AD_ = *R^2^* - *R^2^*_1c_ = 2%) (**Fig. 3B**). Further, age and plasma pTau_181_ explained shared variance in neural selectivity (*R^2^*_Age&AD_ = *R^2^* - *R^2^* - *R^2^* - *R^2^* = 3.7%), as did age and DAN SME (*R^2^* = *R^2^* - *R^2^*_2b_ - *R^2^*_Age_ - *R^2^* = 1.7%) (**Fig. 3C**). Consistent with their independence, DAN SME and plasma pTau_181_ shared little variance (*R^2^* = *R^2^* - *R^2^* - *R^2^* - *R^2^* = 0.1%; **Fig. 3C**).

### Neural selectivity predicts differences in mnemonic function

We next asked if neural selectivity at encoding explains individual differences in within-task and out-of-task cognitive performance, focusing first on selectivity in place-selective cortex on remembered trials, and controlling for age, sex, and education. Greater neural selectivity was associated with high associative *d’* (*β* = .259, *p*_Holm_ = .028; **Fig. 4A**, left), as well as place (*β* = .366, *p*_Holm_ < .001) and face (*β* = .233, *p*_Holm_ = .023) associative *d’* (***Supplementary Results***). Moreover, direct comparison revealed a significantly stronger relationship between associative *d’* and selectivity on remembered than on forgotten trials (*Δr*_Rem-Forg_ = 0.337, 95% CI = [.165, .50]; **Fig. 4A**, left).

**Fig. 4.**
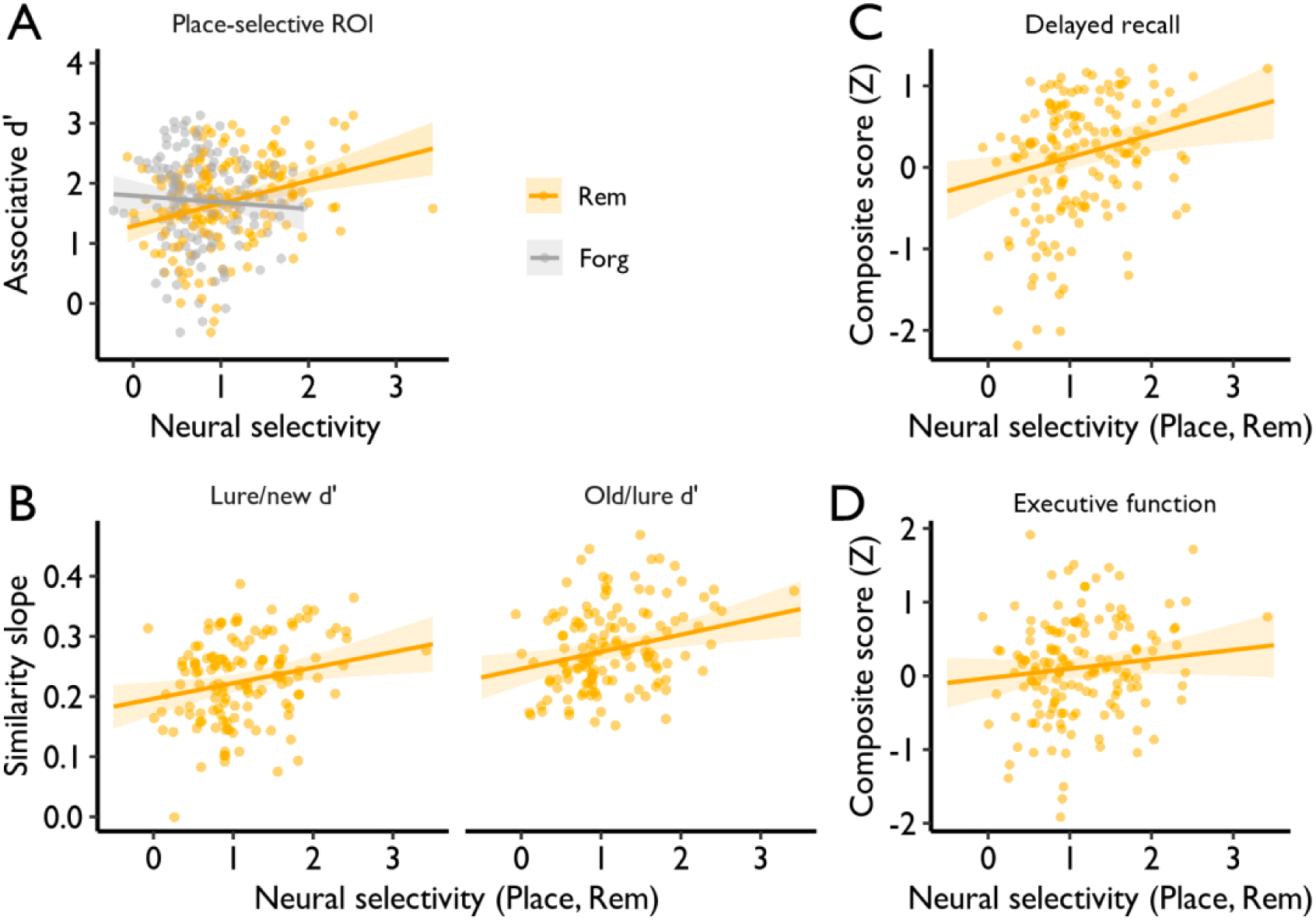
Greater neural selectivity is related to better memory performance. (A) Neural selectivity in place-selective regions predicted associative memory (overall associative *d’*) on remembered trials. (B) & (C) Neural selectivity in place-selective regions (i.e., Place) on remembered trials (i.e., Rem) predicts out-of-task memory performance (B, mnemonic similarity task: the similarity slope on the y-axis reflects the magnitude of the increase in performance as target–lure similarity moved from high to low; C, delayed recall). (D) Neural selectivity in place-selective regions on subsequently remembered trials was not significantly associated with executive function.

Critically, we observed similar effects when using a mnemonic similarity task (MST; see ***Methods***) collected on a different day separated on average by ∼4 weeks from the fMRI session (**Fig. 4B and *Supplementary Results***), suggesting that the relationship between neural selectivity and memory was generalized beyond fMRI task-related memory performance. Similarly, greater neural selectivity was associated with a higher composite delayed recall score collected on average ∼5 weeks from the fMRI session (*β* = .276, *p* = .006; **Fig. 4C**), whereas selectivity was not significantly associated with a composite executive function score (*β* = .128, *p* = .167; **Fig. 4D**), with trend-level evidence that neural selectivity differentially predicted individual differences in memory relative to executive function (Δ_delayed recall – executive function_ = .196, *p* = .062; n = 155). Neural selectivity in face-selective regions on remembered trials did not exhibit significant associations with memory performance (***Supplementary Results***). Collectively, these data indicate that neural selectivity in place-selective cortex partially explains individual differences in both within- and out-of-task memory behavior.

### Multiple pathways account for memory variability in older adults

Considering the significant link between neural selectivity at encoding and individual differences in memory, along with the three factors –– age, AD biomarkers, and top-down attention –– that explain variability in neural selectivity, we developed a SEM to understand pathways from age to neural selectivity to memory. In addition to a pathway from age to neural selectivity to memory, the SEM included two additional pathways through neural selectivity: 1) age → plasma pTau_181_ → neural selectivity → memory performance (AD-related pathway), which tested whether age-related memory decline is explained, in part, by reduced neural selectivity, which, in turn, is explained, in part, by elevated plasma pTau_181_; and 2) age → DAN SME → neural selectivity → memory performance (attention-related pathway), which tested whether age-related decline in top-down attention (DAN SME) explains, in part, reduced neural selectivity, which, in turn, explains, in part, variance in memory (**Fig. 5A**; see ***Methods***). The model included data from 137 participants who had available data for all five factors. Paths not including AD biomarkers were additionally examined in the full sample and replicated the findings in this subsample with all five factors (see ***Supplementary Results***). To simplify the results, we report paths to place associative *d’* below, which is most related to neural selectivity in place-selective regions on remembered trials (see ***Supplementary Results*** for analyses of paths to overall associative *d’* and face associative *d’*; **Table S12**).

**Fig. 5.**
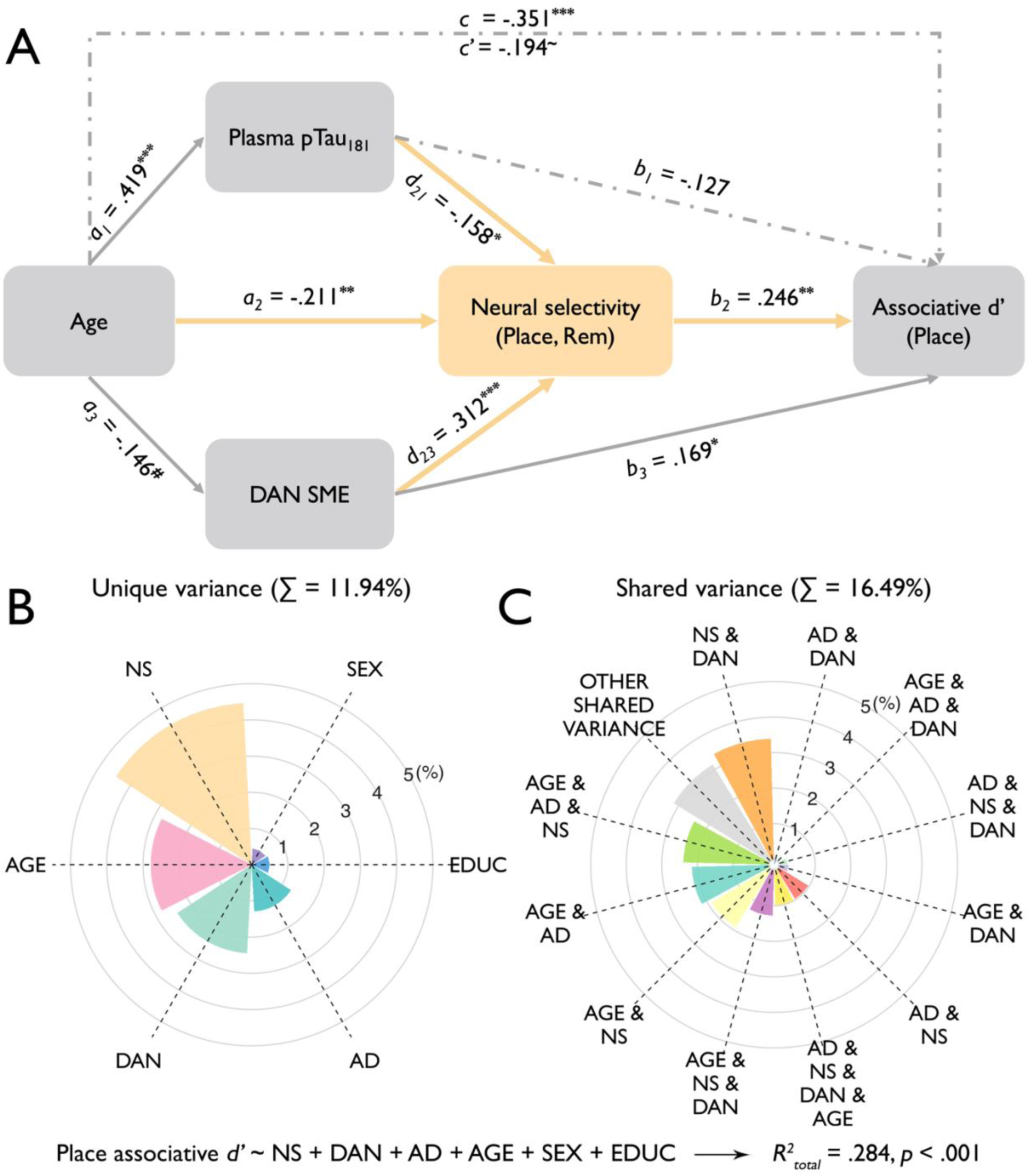
Pathways and predictors on memory performance (i.e., place associative *d’*). (A) Structural equation modelling. Solid lines represent significant paths; dashed lines represent nonsignificant paths. The figure shows standardized betas. **^#^** *p* = .05, **\*** *p* < .05, **\*\*** *p* < .01, **\*\*\*** *p* < .001. *c’* = direct effect; *c* = total effect = *a_1_b_1_ + a_2_b_2_ + a_3_b_3_ + a_1_d_21_b_2_ + a_3_d_23_b_2_ + c’*. (B) Unique and (C) shared explained variances of predictors of place associative *d’*. NS = neural selectivity, DAN = DAN SME (i.e., subsequent memory effect in dorsal attention network), AD = plasma pTau_181_, EDUC = years of education.

First, the SEM revealed that neural selectivity mediated the relationships between age (*a_2_b_2_* = -.006, *β*_standardized_ = -.052, 95% CI = [-.014, -.002]), plasma pTau_181_ (*d_21_b_2_* = -.210, *β*_standardized_ = -.039, 95% CI = [-.506, -.049]), and top-down attention (*d_23_b_2_* = .161, *β*_standardized_ = .077, 95% CI = [.065, .326]) with place associative *d’*. These mediation effects were replicated in separate mediation analyses with our full sample for age (**Table S13**) and top-down attention (**Table S14**). These findings demonstrate that, in CU older adults, neural selectivity plays an important role in associative memory differences that occur with age, diminished top-down attention, and preclinical AD pathology. Second, the SEM revealed that both the AD-(*a_1_d_21_b_2_* = -.002, *β*_standardized_ = -.016, 95% CI = [-.005, -.0005]) and attention-related (*a_3_d_23_b_2_* = -.001, *β*_standardized_ = -.011, 95% CI = [-.004, -.0001]) pathways explain unique variance in memory performance through unique variance in reduced neural selectivity (**Fig. 5A**). Corresponding SEMs predicting out-of-task performance (i.e., delayed recall and MST performance) provided further evidence for multiple pathways between age, neural selectivity, and memory (see ***Supplemental Results***).

To further summarize the unique and shared contributions of age, neural selectivity, DAN SME, plasma pTau_181_, sex, and education in predicting place associative *d’*, we computed hierarchical linear models, as outlined in **Table S11**. The results revealed that neural selectivity explained the largest unique variance in place associative *d’* (*R^2^* = *R^2^* - *R^2^* = 4.4%), followed by age (*R^2^*_Age_ = *R^2^* - *R^2^* = 2.8%) and DAN SME (*R^2^*_DAN_ = *R^2^* - *R^2^* = 2.4%), and finally by plasma pTau_181_ (*R^2^* = *R^2^* – *R^2^* = 1.3%) (**Fig. 5B**). Plasma pTau_181_ shared considerable variance (*R^2^*_sharedAD_ = *R^2^* – *R^2^*= 7.2%) with other predictors, especially with age (*R^2^*_Age&AD_ = *R^2^* – *R^2^* – *R^2^* – *R^2^* = 2.1%), in explaining memory (**Fig. 5C**).

## Discussion

The current study systematically examined mechanisms of intra- and inter-individual differences in neural selectivity and explored their impact on episodic memory in CU older adults. Intra-individual, neural selectivity in place- and face-selective ROIs and activity in frontoparietal nodes of the DAN were significantly higher when encoding later remembered than later forgotten trials. Inter-individual, neural selectivity significantly decreased with age, principally on later remembered trials, with the decrease reflecting attenuated activation to preferred stimuli. Moreover, higher memory-related DAN activity and lower plasma pTau_181_ levels were associated with greater neural selectivity, principally on subsequently remembered trials in place-selective regions. A path analysis integrated the effects of age, DAN activity, plasma pTau_181_, and neural selectivity on memory performance, uncovering distinct age-related and age-independent pathways from AD pathology and diminished top-down attention to reduced neural selectivity that ultimately predicted reduced episodic memory. Collectively, these findings indicate that the quality of cortical representations during learning is influenced by multiple factors, including age, top-down attention, and AD pathology, with distinct interactions between these factors and neural selectivity having an impact on successful episodic memory encoding and later remembering in CU older adults.

Age-related changes in neural selectivity are well documented (*11*–*13, 15, 67*), with prior studies typically adopting a between-groups approach that revealed lower neural selectivity in older compared to younger adults (for individual-difference data, see Park et al (*66*); cf. Voss et al. (*68*)). Here, we used a combined intra- and inter-individual difference approach with a large sample of CU older adults and demonstrate for the first time, to our knowledge, an interaction between age and subsequent memory in neural selectivity. Specifically, neural selectivity during encoding in CU older adults was greater on subsequently remembered compared to forgotten trials, suggesting that, within individuals, the precision of cortical representations of events during learning is important for later remembering (*13, 69*), and moreover, between individuals, this precision on subsequently remembered trials declined with increasing age later in the lifespan. These findings link event-level neural selectivity to event-level later remembering in older adults, suggesting that neural selectivity reflects, in part, the strength or precision of internal (cortical) representations of event features as events unfold and that this precision predicts future remembering when those features are the targets of future retrieval attempts. These findings extend past studies in younger adults that demonstrate that the strength of cortical representations at encoding predict subsequent memory (*24, 25, 70*–*72*), revealing that age-related declines in the precision or strength of cortical representations (*73*) that occur across the later lifespan are differentially observed on subsequently remembered trials. This suggests that even on successfully encoded trials (i.e., events later remembered), memory strength and/or precision continue to decline from one’s 60s to 80s, a possibility that should be explored in future fMRI studies that include quantitative behavioral assays of memory precision (*74*–*77*).

We observed a trend for a greater age-related decline in scene-vs. face-associative memory performance, which was accompanied by evidence that the age-related decline in neural selectivity was greater in scene- than in face-selective regions (see also, Srokova et al. (*78*)). This decline stemmed from reduced activity in response to preferred stimuli (i.e., scenes) rather than an enhanced response to non-preferred stimuli (i.e., faces), consistent with neural attenuation (*66, 78, 79*). Extant data indicate that age-related declines in neural selectivity are consistently observed in scene-selective areas, whereas evidence for changes in neural selectivity in face- and/or object-selective regions is more variable (*11, 16, 68, 78*–*82*). One possibility is that faces consist of more typical (i.e., overlapping) features and, when using famous faces (as in the present study), participants may also shift encoding to differentially rely on semantic vs. perceptual features. By contrast, the visual representations of scenes (including landmarks, as here) may be more distinctive, providing an opportunity to detect age-related variation in perceptual encoding and neural selectivity (*78, 83, 84*). Another possibility is the "last in, first out" hypothesis, which posits that brain regions that mature last are the first to degenerate with age (*3, 85*). While face processing areas appear to mature early (*86, 87*), regions involved in scene processing, especially the RSC and OPA (*88*), appear to emerge later and may reach maturity as late as early adulthood (*89*–*91*). This delayed maturation may render scene-selective regions more susceptible to factors that drive age-related declines in neural selectivity.

The present focus on factors driving age-related declines in neural selectivity revealed that top-down attention during encoding (a) predicts subsequent memory (*28, 30, 65, 92*), (b) modulates neural selectivity (*17, 22, 26*), (c) does not vary with plasma or CSF biomarkers of preclinical AD, and (d) partially accounts for age-independent and age-related variability in memory through its impact on neural selectivity. Controlling for age, CU older adults with greater memory-related neural activity in the DAN demonstrated higher neural selectivity during subsequently remembered events, with the DAN SME being the top predictor of variance in neural selectivity. Moreover, this age-independent association between neural selectivity and DAN activity was significantly stronger compared to neural selectivity’s relationship with VAN activity, indicating that neural selectivity during successful encoding is predominantly modulated by top-down rather than bottom-up attention (*54, 55, 93, 94*). The SEM also revealed that age-related differences in memory stem, in part, from age-related change in DAN activity, which leads to declines in neural selectivity.

Potentially in line with our findings, prefrontal dopamine D1-receptor-mediated activity enhances the magnitude and orientation selectivity of neural responses in the visual cortex of macaques (*95*); prefrontal dopamine plays a central role in the modulation of top-down attention (*96, 97*); and, with age, the densities of dopamine D1/D2, serotonin, and acetylcholine receptors in human frontal cortex significantly decrease, contributing to age-related cognitive dysfunction (*98*). Aging is also associated with structural (*58*–*60*) and functional (*61, 62*) changes in and between frontoparietal nodes of the DAN, which likely contribute to disruption of top-down attention. Thus, attention-mediated age-related decline in neural selectivity may result, in part, from dysregulated frontoparietal neurotransmission –– possibly stemming from receptor, neurotransmitter, and connectivity changes –– that disrupts top-down attentional modulation of perceptual representations of event features. The present observation that reduced neural selectivity reflects attenuated activation to preferred stimuli is compatible with diminished top-down gain modulation.

While early AD pathology (e.g., plasma pTau_181_) was unrelated to memory-related engagement of top-down attention, our data reveal that early AD pathology was associated with reduced neural selectivity and partially accounted for age-related memory decline via neural selectivity. Moreover, we observed an age-independent relationship between early AD pathology and neural selectivity. Consistent to previous tau PET findings (*48*), neural selectivity’s negative relationship with early AD pathology was principally observed in place-selective regions. However, whereas Maass et al. (2019) (*48*) observed evidence for functional response broadening (i.e., increased activation to non-preferred categories), the present findings are ambiguous with respect to whether neural selectivity change with early AD pathology reflects attenuation and/or broadening (**Fig. 2F**). Given that our study only examined global biofluid measures of AD (plasma and CSF measures of pTau_181_), we are unable to address whether differences in neural selectivity relate to focal tau within the medial temporal lobe and/or cortical regions important for category-specific effects.

The present data revealed that neural selectivity accounted for age-independent variability in memory, not only within-task (i.e., fMRI associative memory performance) (*16, 78, 79*) but also across-task (i.e., MST and delay-recall composite performance measured weeks apart from the fMRI session). In contrast, neural selectivity did not vary with a composite executive function measure that spanned processing speed (*79*), working memory, and verbal fluency (*78*). Evidence that neural selectivity at encoding predicts performance on multiple memory measures acquired in different temporal and spatial contexts implies that individual differences in selectivity relate, in part, to stable differences in memory abilities (i.e., is “trait”-like), rather than solely to transient state differences between people at the time of measurement. This observation should motivate future research to explore whether neural selectivity predicts longitudinal memory decline and progression to mild cognitive impairment.

Related to the latter, given the cross-sectional nature of the present study, it is difficult to establish causality and temporal relationships between neural selectivity and other variables. An ongoing longitudinal study using a 7-year follow-up with the present sample will ultimately help address these questions. We also note that, although the present sample size of deeply phenotyped older adults is relatively large, it lacks educational and racial/ethnic diversity which potentially limits the generalizability of the findings. Future studies should include more diverse samples, which is also one of our future directions. Finally, while we observed that early AD biomarkers relate to neural selectivity and memory performance, but not to top-down attention, the variance explained by these biomarkers was modest, especially in the subsample of participants with available CSF. This likely reflects our focus on CU older adults, who by definition are characterized as demonstrating cognition (including memory) in the normal range, along with the restricted number of participants with elevated AD biomarkers. PET assays of regional amyloid and tau burden may ultimately provide sensitive and complementary insights into how AD pathology impacts neural selectivity and, in doing so, memory.

In sum, this study revealed age-related and age-independent factors that influence cortical representations during event encoding and predict subsequent memory performance. Integrating age, early AD pathology, top-down attention, and neural selectivity, we elucidated multiple pathways underlying individual differences in episodic memory among CU older adults. On the one hand, controlling for age, AD-independent reductions in top-down attention and AD-related pathology alter the precision of cortical representations of event features during experience, with consequences for future remembering. On the other hand, age-related episodic memory decline is partially accounted for by reduced neural selectivity, that is in turn independently impacted by top-down attention and early AD pathology. As such, these data inform models of the factors –– related to and independent of age and related to and independent of early AD processes –– that impact memory formation and account for why some CU older adults remember better than others.

## Materials and methods

### Participants

This study includes data from 166 CU older adults (**Table 1**; 60–88 years, 95 female) of an initial 212 participants (60–88 years, 121 female) enrolled in the SAMS(*8, 37*). SAMS eligibility included: normal or corrected-to-normal vision and hearing; right-handed; native English speaking; no history of neurological or psychiatric disease; a Clinical Dementia Rating score of zero (CDR)(*99*) and performance within the normal range on a neuropsychological assessment. Inclusion in the current analyses additionally required the availability of both structural MRI and task-based fMRI data, which were collected from 170 participants. Among these, 4 participants were excluded from all analyses due to excess head motion during scanning (n = 3; see fMRI pre-processing) or visible artifacts in the fMRI data (n = 1). This resulted in available data from 166 older adults. Of the 166 participants with an acceptable quality of fMRI data, 10 participants were excluded from the individual-difference analyses due to having fewer than 3 trials in at least one of the four conditions [category (i.e., face or place) × memory (i.e., remembered vs forgotten)]. Thus, 156 participants are included in the individual-differences analyses (**Table 1**). All participants provided informed consent in accordance with a protocol approved by the Stanford Institutional Review Board.

**Table 1.**
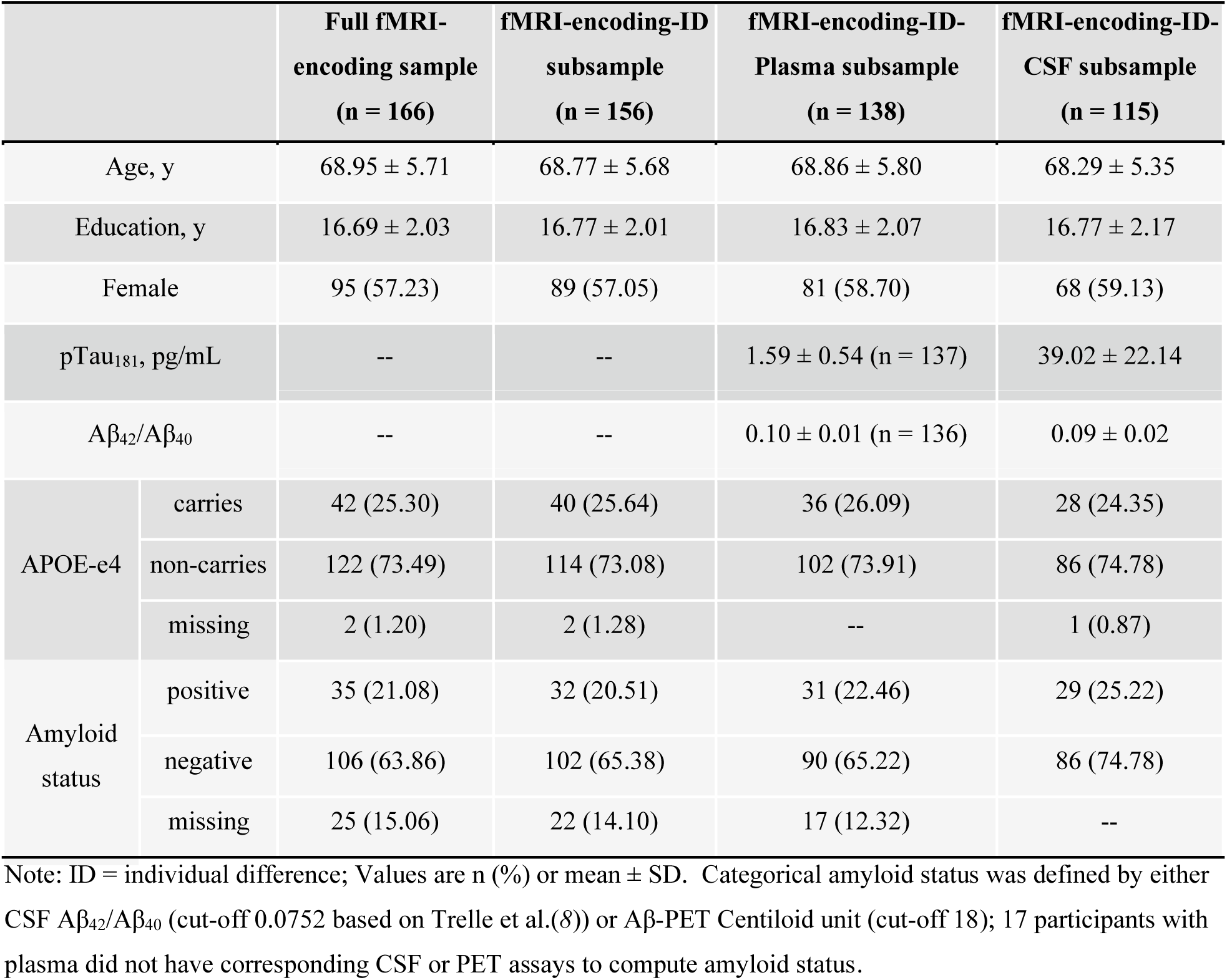
Participant Demographics and Biomarker Summary.

|  |  | Full fMRI-encoding sample<br>(n = 166) | fMRI-encoding-ID<br>subsample<br>(n = 156) | fMRI-encoding-ID-Plasma subsample<br>(n = 138) | fMRI-encoding-ID-CSF subsample<br>(n = 115) |
| --- | --- | --- | --- | --- | --- |
| Age, y |  | 68.95 ± 5.71 | 68.77 ± 5.68 | 68.86 ± 5.80 | 68.29 ± 5.35 |
| Education, y |  | 16.69 ± 2.03 | 16.77 ± 2.01 | 16.83 ± 2.07 | 16.77 ± 2.17 |
| Female |  | 95 (57.23) | 89 (57.05) | 81 (58.70) | 68 (59.13) |
| pTau<sub>181</sub>, pg/mL |  | -- | -- | 1.59 ± 0.54 (n = 137) | 39.02 ± 22.14 |
| Aβ<sub>42</sub>/Aβ<sub>40</sub> |  | -- | -- | 0.10 ± 0.01 (n = 136) | 0.09 ± 0.02 |
| APOE-e4 | carries | 42 (25.30) | 40 (25.64) | 36 (26.09) | 28 (24.35) |
|  | non-carries | 122 (73.49) | 114 (73.08) | 102 (73.91) | 86 (74.78) |
|  | missing | 2 (1.20) | 2 (1.28) | -- | 1 (0.87) |
| Amyloid<br>status | positive | 35 (21.08) | 32 (20.51) | 31 (22.46) | 29 (25.22) |
|  | negative | 106 (63.86) | 102 (65.38) | 90 (65.22) | 86 (74.78) |
|  | missing | 25 (15.06) | 22 (14.10) | 17 (12.32) | -- |

Note: ID = individual difference; Values are n (%) or mean ± SD. Categorical amyloid status was defined by either CSF Aβ<sub>42</sub>/Aβ<sub>40</sub> (cut-off 0.0752 based on Trelle et al.(8)) or Aβ-PET Centiloid unit (cut-off 18); 17 participants with plasma did not have corresponding CSF or PET assays to compute amyloid status.

### Biofluid Data

#### Plasma data

138 participants completed a blood draw during the study enrollment, which preceded the fMRI session (mean lag = 5.28 weeks; SD = 3.38 weeks). EDTA plasma was collected by venipuncture, centrifuged for 10 minutes at 2000 x g at 22°C, aliquoted in polypropylene tubes and stored at -80°C until measurement. Using procedures previously described(*100*) , plasma Aβ_42_ and Aβ_40_ (n = 136 participants) and pTau_181_ (n = 137 participants) were measured using the fully automated Lumipulse *G* 1200 instrument (Fujirebio US, Inc.,Malvern, PA). A single aliquot for each participant was used to measure Aβ_42_ and Aβ_40_ in a single batch analysis, and a single aliquot was used to measure pTau_181_ in a separate single batch analysis. Analyses were conducted by the Stanford Alzheimer’s Disease Research Center (ADRC) Biomarker Core.

#### CSF data

115 participants completed a CSF draw within a mean of 4.23 weeks (SD = 4.81 weeks) of the fMRI session. As described previously(*8*), CSF samples were centrifuged for 15 minutes at 500 x g at 22°C, aliquoted in polypropylene tubes and stored at -80°C until measurement. Using procedures previously described(*101*), a single aliquot for each participant was used to measure Aβ_42_, Aβ_40_, pTau_181_, and total tau using the fully automated Lumipulse *G* 1200 instrument in a single-batch analysis by the Stanford ADRC Biomarker Core.

### In-scanner Task

#### Word-picture associative memory task

The word-picture associative memory task (**Fig. 1A**) was administered concurrent with fMRI as previously described(*8, 37*). Briefly, participants first encoded word-face and word-place associations and then engaged in an associative retrieval task. The word–picture pairs comprised concrete nouns (e.g., “banana,” “violin”) paired with pictures of famous faces (e.g., “Queen Elizabeth,” “Ronald Reagan”) or well-known places (e.g., “Golden Gate Bridge,” “Niagara Falls”). The task consisted of 5 alternating study and test blocks. Each study block included 12 word–face and 12 word–place pairs; participants were instructed to form a link between the word and picture presented. In each test block, participants saw a mix of 24 studied words and 6 novel (foil) words. Memory was assessed using an associative cued-recall test accompanied by a button response: participants selected “face” or “place” if they remembered the word and could recall the associated picture or picture category; they selected “old” if they remembered the word but could not recall the associated picture or picture category; they selected “new” if they did not remember studying the word. Associative memory performance was estimated using a sensitivity index, associative *d′*, where hits were defined as correct associative category responses to studied words and false alarms were defined as incorrect category responses to new words. Thus, associative *d′* = Z(“correct associate category”|old) − Z(“associate category”|new). Similar calculations were performed for place [Z(“place correct associate category”|old place-word pairs) − Z(“place associate category”|new)] and face [Z(“face correct associate category”|old face-word pairs) − Z(“face associate category”|new)] associative *d’*, respectively.

#### MRI Data Acquisition and Preprocessing

Data were acquired on a 3T GE Discovery MR750 MRI scanner (GE Healthcare) using a 32-channel radiofrequency receive-only head coil (Nova Medical). Functional data were acquired using a multiband EPI sequence (acceleration factor = 3) consisting of 63 oblique axial slices parallel to the long axis of the hippocampus (TR = 2 s, TE = 30 ms, FoV = 215 mm × 215 mm, flip angle = 74°, voxel size = 1.8 × 1.8 × 2 mm). To correct for B0 field distortions, we collected two B0 field maps before every functional run, one in each phase encoding direction. Structural MRI included a whole-brain high-resolution T1-weighted anatomical volume (TR = 7.26 ms, FoV = 230 mm × 230 mm, voxel size = 0.9 × 0.9 × 0.9 mm, slices = 186).

MR data were preprocessed using fMRIPrep 23.0.0rc0(*102*) (RRID:SCR_016216), which is based on Nipype 1.8.5(*103, 104*) (RRID:SCR_002502). Functional images were corrected for susceptibility distortion, head motion, and slice-timing, and co-registered to the T1w structural volume. For the group-level analysis, we resampled BOLD runs into standard volumetric (i.e., *ICBM 152 Nonlinear Asymmetrical template version 2009c*) and surface (i.e., *fsaverage*) spaces. Grayordinates files(*105*) containing 91k samples were generated using the highest-resolution *fsaverage* as intermediate standardized surface space.

Images with motion artifacts were automatically identified as those TRs in which total displacement relative to the previous frame exceeded 0.9 mm. Trials with framewise displacement (FD) of any timepoints exceeding 0.9 mm were identified as artifacts. Runs in which the number of artifacts identified exceeded 25% of TRs or 50% of trials, as well as runs in which FD exceeded 5 mm, were excluded. These criteria led to exclusion of data from three participants who exhibited excess head motion across runs, as well as exclusion of a varying number of study and test runs from five additional participants. Both encoding and retrieval fMRI data were preprocessed, but we report only on encoding data in the current study (for results at retrieval, see Trelle et al.(*37*)).

### Whole-Brain Univariate Analysis

Before conducting univariate activation analyses, surface-based and volume-based data were spatially smoothed using a 6-mm FWHM Gaussian kernel and filtered in the temporal domain using a nonlinear high-pass filter with a 100-s cutoff. Specifically, the *wb_command* was used for surface-based smoothing and *SUSAN* was used for volume-based smoothing. Univariate activation analysis in volumetric space was performed for a leave-one-subject-out procedure (see *Definition of ROIs;* **Fig. S2**) and ROI activity extraction (see *ROI-based Univariate Analysis*). General linear modeling within the FILM module of FSL was used to model the data. During encoding, word-face and word-place pairs were conditioned on subsequent memory, modeling subsequent associative hits, associative misses, item hits, and item misses separately. Events were modeled as boxcar functions based on the time of stimulus onset and duration (4s) and convolved with the canonical hemodynamic response function (double gamma function). The category effect was defined as the difference between word-face and word-place pairs, and the SME was defined as the difference between subsequently remembered associations (i.e., associative hits) and forgotten associations (i.e., associative misses, item hits, and item misses). These contrasts were then submitted to group-level analyses by using FSL Permutation Analysis of Linear Models (PALM; https://fsl.fmrib.ox.ac.uk/fsl/fslwiki/PALM) with 1,000 iterations and tail acceleration(*106*). Unless otherwise noted, significant results are reported after whole-brain correction for multiple comparisons, using a two-step approach: first, controlling for family-wise error (FWE) within each of the left hemisphere, right hemisphere, and subcortical regions with p < 0.05 using threshold-free cluster enhancement (TFCE)(*107*), and second, using Bonferroni correction for the three macroscopic regions (i.e., left hemisphere, right hemispheres, and subcortical; i.e., -log10(alpha/N) = -log10(0.05/3) = 1.7782).

### Definition of ROIs

The primary neural selectivity analyses were conducted in face- and place-selective cortical ROIs defined in a two-step process. First, a priori ROIs were selected based on an independent probabilistic functional atlas that shows the likelihood of a given voxel being located in the face- or place-selective regions (probability > 0.1)(*108*), with the OFA and FFA defined as face-selective regions, and the OPA, PPA, and RSC defined as place-selective regions (**Fig. 1D**). Second, these masks were intersected with whole-brain univariate category effect masks from the current experiment’s encoding phase data. To ensure statistical independence, we used a leave-one-participant-out procedure to define ROIs for each held-out participant (**Fig. S2**). Specifically, for a given participant, we performed group-level analysis excluding this participant and obtained a whole-brain category effect map in MNI space, after controlling for FWE with *p* < 0.05 through TFCE(*107*). We then intersected the participant-independent whole-brain category map with the predefined functional ROIs(*108*), and each resulting ROI mask was transformed to the participant’s native space using *antsApplyTransforms* and defined as all voxels intersecting the midpoint between the gray-white and gray-pial boundaries. For the OFA ROI, 10 participants were excluded from the analysis (number of voxels < 30) due to having only partial coverage of early visual cortex in their field of view.

We additionally examined the role of attention in modulating neural selectivity during encoding. To address this question, we used the Schaefer 2018 atlas (7-network with 400 parcels)(*109*) to define frontoparietal nodes of the DAN, which included the frontal eye fields (FEF) and intraparietal sulcus (IPS), and the VAN, which included the temporoparietal junction (TPJ) and the inferior frontal cortex (IFC). Specifically, FEF contained all FEF nodes within the DAN (ROI labels: 86-89 for left and 290-292 for right), IPS contained parietal nodes within the DAN (ROI labels: 74-85 for left and 276-289 for right), TPJ contained temporal occipital and temporal parietal nodes within the VAN (ROI labels: 92-95 for left and 296-300 for right), and IFC contained frontal-operculum-insula nodes within the VAN (ROI labels: 97-105 for left and 302-309 for right). The DAN and VAN ROIs were transformed to the participant’s native space and intersected with the midpoint between the gray-white and gray-pial boundaries.

### ROI-based Univariate Analysis

All ROI-based analyses were conducted in the participants’ native space (i.e., T1 space). To examine neural selectivity in each face- and place-preferred region, we performed a whole-brain subject-level univariate analysis and extracted the mean contrast estimate (t-value) of face vs. place trials from the ROI. Neural selectivity in face-preferred regions (i.e., FFA and OFA) was defined as face – place, and vice versa for place-preferred regions (i.e., PPA, OPA, and RSC). Similarly, we extracted face- and place-related activity (relative to the implicit baseline) in each face- and place-preferred ROI. The univariate SME in the DAN (i.e., mean t-value in FEF and IPS) and VAN (i.e., mean in TPJ and VFC) was extracted from the contrast of remembered trials (i.e., associative hit trials) vs. forgotten trials (i.e., associative miss, item hit, and item miss trials).

### Out-of-scanner Cognitive Tasks

#### Standardized cognitive composite scores

Test scores from a neuropsychological battery completed by all participants during study enrollment(*37*) were used to create two composite cognitive scores: a delayed recall score and an executive function score. The delayed recall score(*8, 37*) was comprised of performance across (1) the logical memory subtest of the Wechsler Memory Scale, (2) the Hopkins Verbal Learning Test–Revised, and (3) the Brief Visuospatial Memory Test–Revised. The executive function score was comprised of performance across tests measuring (1) task switching (Trail Making Test B, inverse), (2) working memory (WMS-III digit span total score), and (3) category fluency (animal naming). Composite scores were computed by first z-scoring individual subtest scores using the full SAMS sample as a reference and then averaging. These neuropsychological data were collected on a separate day and preceded fMRI data collection by a mean of 4.89 weeks (SD = 3.76 weeks).

#### MST

The MST was administered using previously described measures and instructions(*8, 110*) and was collected within a mean of 4.30 weeks (SD = 4.04 weeks) of the fMRI session. During an incidental encoding phase, participants made indoor/outdoor judgments for 128 pictures of everyday objects. Participants then performed a surprise memory test, in which 64 of the studied objects were intermixed with 64 perceptually similar lure objects and 64 novel (dissimilar) objects. Participants were to respond “old” if they remembered the object as having been studied, “similar” if they remembered the object as similar, but not identical, to a studied object, or “new” if they remembered the object as not having been studied. The 64 lures systematically varied in perceptual similarity to the untested studied objects across 5 levels of similarity/difficulty. Here performance was estimated for each level of target-lure similarity using two sensitivity indices: (1) lure/new d’, the ability to correctly classify perceptually similar lures and differentiate them from novel objects, as Z(“similar” | lure) − Z(“similar” | novel foil), and (2) old/lure *d′*, the ability to correctly endorse studied objects and avoid the propensity to incorrectly endorse lures as old, as Z(“old” | target) − Z(“old” | lure). Building on prior work(*8*), the primary measure of interest was the degree to which performance improved as a function of decreasing target-lure similarity (i.e., decreasing task difficulty). LMMs were used to model performance across the 5 similarity levels and to examine interactions with measures of neural selectivity. LMMs included a random intercept and slope for each participant. Participant-specific slopes, which reflected the magnitude of the increase in performance as target-lure similarity moved from high to low (i.e., decreasing difficult) was extracted and plotted against neural selectivity to visualize similarity × selectivity interactions. MST data were available from 135 participants.

### Statistical Analysis

#### LMM

For repeated-measures models exploring effects of category (face, place), subsequent memory (remembered, forgotten), and region (face-selective, place-selective) on neural selectivity, LMMs were used to examine the relationship between dependent and independent variables with *lmer* function from the *lmerTest* package(*111*) in R 4.3.0 (R Core Team 2023). Participants were set as random effects. The degrees of freedom in the mixed models was estimated by the Satterthwaite approximation(*111*). Unless otherwise specified, *p* values were corrected using Holm-Bonferroni correction for multiple comparisons (termed as *p*_Holm_ in the text).

#### Direct comparisons for two correlation coefficients

To examine whether neural selectivity on subsequently remembered trials was more strongly related to memory performance than on forgotten trials, we conducted direct comparison analyses using the R package ‘cocor’(*112*), which provides a comprehensive solution to compare two correlations based on dependent groups with overlapping variables (i.e., associative *d’*). We report statistical results in the main text based on the computation of confidence intervals (CI)(*113*) since such tests are generally considered superior to significance tests because they indicate, respectively, the magnitude and precision of the estimated effect(*114, 115*).

#### Mediation analysis

Mediation effect tests were implemented with R package *lavaan*(*116*). For simple mediation analyses, we examined the relationship between (1) mediators (M) and independent variables (X) (M = k_1_ + *a*X + *ε_1_*); and (2) independent variables (X) and dependent variables (Y) with mediator (M) (Y = k_2_ + *c’*X + *b*M + *ε_2_*). Sex and years of education were included in models as confounding variables. We also included age as a covariate when age was not a variable of interest in the model. The indirect effect was estimated as *ab* and the direct effect was estimated as *c’*. The direct and indirect effects sum to the total effect, *ab+c’*.

A multiple mediation analysis was used to examine the mediation effects of early AD pathology (plasma pTau_181_) and top-down attention (DAN SME) for the relationship between age and neural selectivity. Specifically, we examined the relationships between (1) plasma pTau_181_ (M1) and age (X) (M1 = k_1_ + *a_1_*X + *ε_1_*); (2) top-down attention (M2) and age (X) (M2 = *a_2_*X + *ε_2_*); and (3) neural selectivity (Y) with age (X), plasma pTau_181_ (M1), and top-down attention (M2) (Y = k_2_ + *c’*X + *b_1_*M1 + *b_2_*M2 + *ε_3_*). Sex and years of education were included in the model as confounding variables. In the above equations, X (age) is the predictor, Y (neural selectivity in place-selective region on later remembered trials) is the dependent variable, and M1 (plasma pTau_181_) and M2 (DAN SME) are the mediators.

Unless otherwise specified, the significance of indirect effects was determined by 95% CI based on 5,000 bootstrap resampling. In addition, to ensure comparability of *β* values across different pathways, we also report standardized *β* estimates using a completely standardized solution (i.e., *std.all*) in the *parameterEstimates* function provided by *lavaan*(*116*). The same approaches were used for the SEM below.

#### SEM

The *lavaan*(*116*) was used to conduct SEM to examine whether individual differences in episodic memory in CU older adults were impacted by two independent pathways (**Fig. 5A**): (1) age → plasma pTau_181_ → neural selectivity (restricted to place-selective regions on later remembered trials, hereafter) → memory performance, and (2) age → top-down attention → neural selectivity → memory performance. To do this, we tested the relationships between (1) plasma pTau_181_ (M1) with age (X) (M1 = k_1_ + *a_1_*X + *ε_1_*); (2) neural selectivity (M2) with age (X), top-down attention (M3; i.e., DAN SME), and plasma pTau_181_ (M1) (M2 = k_2_ + *a_2_*X + *d_21_*M1 + *d_23_*M3 + *ε_2_*); (3) top-down attention (M3) with age (X) (M3 = *a_3_*X + *ε_3_*); and (4) memory performance (Y) with age (X), plasma pTau_181_ (M1), neural selectivity (M2), and top-down attention (M3) (Y = k_3_ + *c’*X + *b_1_*M1 + *b_2_*M2 + *b_3_*M3 + *ε_4_*). Sex and years of education were included to the model as confounding variables. In the above equations, X (age) is the predictor, Y (memory performance, e.g., associative d’, delayed recall composite score, and MST old/lure d’ similarity slope) is the dependent variable, and M1 (plasma pTau_181_), M2 (neural selectivity in place-selective region on later remembered trials), and M3 (DAN SME) are the mediators. The indirect effect of age → plasma pTau_181_→ neural selectivity → memory performance was estimated as *a_1_×d_21_×b_2_*, and the other indirect effect of age → top-down attention → neural selectivity → memory performance was estimated as *a_3_×d_23_×b_2_*. The total effect (c) equals *c’+a_1_d_21_b_2_+a_1_b_1_+a_2_b_2_+a_3_b_3_+a_3_d_23_b_2_*.

## Classification

Social Sciences/Psychological and Cognitive Neurosciences

## Author Contributions

A.D.W. conceived of the project and designed the experiments; J.S., A.D.W., A.N.T., and E.C.M. conceptualized the study; A.N.T. programmed the fMRI experiment; A.N.T., A.R., J.P., T.T.T., S.J.S., and K.I.A. performed the experiments and collected data; J.S., A.N.T., E.N.W., and T.T.T. analyzed the data; J.S. visualized the results and wrote the original draft; A.D.W., A.N.T., E.C.M., and J.S. reviewed and edited the manuscript. All authors approved the final version of the manuscript for submission.

## Acknowledgments

We thank all members of the Wagner and Mormino Laboratories for their discussions and support. We are grateful to: Adam Kerr, Hua Wu, Michael Perry, and Laima Baltusis at Stanford’s Center for Cognitive and Neurobiological Imaging (CNI) for their assistance in fMRI data acquisition; Jeff Bernstein, Clementine Chou, Nicole Corso, Scott Guerin, Wanjia Guo, Marc Harrison, Madison Hunt, Anna Khazenzon, Celia Litovsky, Manasi Jaykumar, Madison Kist, Ayesha Nadiadwala, Austin Salcedo, Natalie Tanner, and Monica Thieu for their assistance with data collection; and the SAMS volunteers for their participation in the study. This research was supported by the National Institutes of Health (National Institute on Aging; R01AG048076, R01AG074339, and R21AG058859 to Dr. Elizabeth Mormino and Dr. Anthony Wagner and K99AG075184 to Dr. Alexandra Trelle) and The Phil and Penny Knight Initiative for Brain Resilience at the Wu Tsai Neurosciences Institute, Stanford University (to Dr. Jintao Sheng and Dr. Edward Wilson).

## Supplementary Results

### Relationship between age and AD biomarkers

Age was associated with increased log-transformed pTau_181_ (plasma: *β* = .009, *p* < .001; CSF: *β* = .013, *p* < .001) and decreased Aβ_42_/Aβ_40_ ratio (plasma: *β* = -.0005, *p* = .026; CSF: *β* = -.001, *p* = .006). The plasma and CSF assays were correlated (pTau_181_: *r* = .37, *p* < .001, n = 104; Aβ_42_/Aβ_40_: *r* = .51, *p* < .001, n = 103).

### Relationship between bottom-up attention and neural selectivity

To examine if attention effects on neural selectivity are specific to DAN-mediated top-down attention or reflect more general impacts of attentional processes on selectivity, we conducted parallel analyses with frontoparietal regions of the ventral attention network (VAN; **Fig. 1E**) linked to bottom-up attentional processes (*1*). While VAN activity demonstrated a significant SME (*t*_155_ = 3.98, *p* < .001), its magnitude did not significantly vary with age (*β* = .001, *p* = .880). A linear mixed-effect model (LMM), with bottom-up attention (VAN SME), memory (Rem vs. Forg), and region (Face-vs. Place-selective ROI) as predictors of neural selectivity, revealed a three-way interaction (*F*_1,462_ = 4.276, *p* = .039), but the main effect of bottom-up attention was not significant (*F*_1,151_ = 1.608, *p* = .207). Moreover, the simple effect between VAN SME and neural selectivity was not significant in place-(Rem: *β* = .302, *p*_Holm_ = .124; Forg: *β* = -.158, *p*_Holm_ = .603) nor face-selective regions (Rem: *β* = .114, *p*_Holm_ = .603; Forg: *β* = .179, *p*_Holm_ = .603) (**Fig. S4B**). Critically, a direct comparison revealed that the association between DAN SME and neural selectivity on remembered trials was significantly stronger than that observed with VAN SME (Place-selective ROI: Δ*r*_DAN-VAN_ = .188, 95% CI = [.084, .292]; Face-selective ROI: Δ*r*_DAN-VAN_ = .141, 95% CI = [.034, .245]). These results provide evidence for a stronger role of top-down, relative to bottom-up, attention in modulating neural selectivity in category-selective cortex during memory encoding in CU older adults.

### Neural selectivity in place-selective regions is associated with face and place associative *d’*

In place-selective regions, we found that neural selectivity related to both place (Rem: β = .366, p_Holm_ < .001; Forg: β = -.323, p_Holm_ = .053; Δr_Rem-Forg_ = 0.503, 95% CI = [.337, .656]) and face associative *d’* (Rem: β = .233, p_Holm_ = .023; Forg: β = -.203, p_Holm_ = .215; Δr_Rem-Forg_ = 0.345, 95% CI = [.174, .507]), with LMMs revealing a significant interaction between memory category (i.e., place vs. face associative *d’*) and neural selectivity on later remembered trials (*F*_1,154_ = 5.616, *p* = .019), but not on forgotten trials (*F*_1,154_ = .416, *p* = .520). This later result suggests that neural selectivity at encoding in place-selective regions differentially contributes to subsequent place associative *d’* than face associative *d’*.

### Neural selectivity in place-selective regions is associated with mnemonic similarity task performance

Neural selectivity in place-selective regions also explained variance in independent (i.e., out-of-task) memory measures collected during a separate study session separated by weeks from the fMRI sessions. Specifically, neural selectivity on remembered trials related to lure discrimination in the mnemonic similarity task (MST), with a significant similarity × selectivity interaction (**Fig. S8**; lure/new *d’*: *F*_1,130_ = 5.570, *p* = .020; old/lure *d’*: *F*_1,130_ = 9.711, *p* = .002) revealing higher selectivity was associated with better MST performance at lower-levels of target-lure similarity (**Fig. S8**); that is, there were larger increases in discrimination performance with decreasing levels of target-lure similarity in individuals with higher neural selectivity relative to those with lower neural selectivity (**Fig. 4C**).

### Relationship between neural selectivity in face-selective regions and memory performance

In face-selective regions, overall associative *d’* did not significantly relate to neural selectivity on remembered trials (*β* = .093, *p*_Holm_ = .367), but negatively related to neural selectivity on forgotten trials (*β* = -.319, *p*_Holm_ = .026) (**Fig. S9**), and their difference was significant (Δr_Rem-Forg_ = 0.293, 95% CI = [.133, .445]). Although the relationship for subsequently remembered trials was not significant in face-selective regions, it did not statistically differ from that in place-selective regions (Δr_Face-Place_ = -.136, 95% CI = [-.339, .071]); a similar result was observed for forgotten trials (Δr_Face-Place_ = -.092, 95% CI = [-.320, .139]). Similar results were observed when focusing on face associative *d’* (Rem: *β* = .086, *p*_Holm_ = .342; Forg: *β* = -.338, *p*_Holm_ = .004; Δr_Rem-Forg_ = 0.341, 95% CI = [.183, .492]) or place associative *d’* (Rem: *β* = .045, *p*_Holm_ = .647; Forg: *β* = -.254, *p*_Holm_ = .053; Δr_Rem-Forg_ = 0.220, 95% CI = [.060, .378]). Mixed-effects models showed that the interaction between memory category (i.e., place vs face associative *d’*) and neural selectivity in face-selective regions was neither significant on remembered trials (*F*_1,154_ = .034, *p* = .853) nor on forgotten trials (*F*_1,154_ = .534, *p* = .466). These results suggest that neural selectivity in face-selective regions was not related to memory performance on remembered trials, but negatively related to both face and place associative *d’* on forgotten trials.

### SEMs for overall associative *d’* and face associative *d’*

In addition to predictors of place associative *d’* (reported in the main text), when SEMs were constructed using overall and face associative *d’*, none of the memory-related pathways were significant (**Table S12**). Nonetheless, with the full sample (n = 156), we observed significant mediation effects from age → neural selectivity → associative memory (**Table S13**) and DAN SME → neural selectivity → associative memory (**Table S14**).

### SEMs for out-of-task performance

We also generated SEMs that explain out-of-task memory performance (delayed recall composite score, n = 137; MST lure/new *d’* and old/lure *d’* similarity slopes, n = 119). The SEMs revealed a significant AD-related pathway (*a_1_d_21_b_2_* = -.0015, 95% CI = [-.004, -.0001]), but nonsignificant attention-related pathway (*a_3_d_23_b_2_* = -.001, 95% CI = [-.003, .0000]), for the delayed recall composite score (**Table S15**). When individual mediation models were performed with the full sample (n = 156), the effects from age → neural selectivity → delayed recall composite score (**Table S16**) and DAN SME → neural selectivity → delayed recall composite score were both significant (**Table S17**). The SEM explaining MST performance (n = 119) revealed that neither the AD-(*a_1_d_21_b_2_*) nor attention-related (*a_3_d_23_b_2_*) pathway was significant, whereas the pathway from DAN SME → neural selectivity → similarity slope of lure/new *d’* (*d_23_b_2_*) was significant (**Table S18**). Similarly, mediation models with a slightly larger subsample (n = 135) showed neural selectivity significantly mediated the relationship between age and MST performance (**Table S19**) and between top-down attention and MST performance (**Table S20**). These findings suggest that there are multiple pathways leading to individual differences in episodic memory in CU older adults, including elevated plasma pTau_181_ with age that is linked to reduced memory through reduced neural selectivity, and reduced top-down attention that reduces episodic memory via decreased neural selectivity.

### Supplementary Methods MRI data preprocessing

Results included in this manuscript come from preprocessing performed using fMRIPrep 23.0.0rc0 (*2*) (RRID:SCR_016216), which is based on Nipype 1.8.5 (*3, 4*) (RRID:SCR_002502). The below is an edited version of text automatically generated by fMRIPrep.

#### Preprocessing of B0 inhomogeneity mappings

A total of two fieldmaps were found available within the input BIDS structure. A B0-nonuniformity map (or fieldmap) was estimated based on two (or more) echo-planar imaging (EPI) references with topup (*5*) (FSL 6.0.5.1:57b01774).

#### Anatomical data preprocessing

One T1-weighted (T1w) images was collected in the current study. The T1w image was corrected for intensity non-uniformity (INU) with N4BiasFieldCorrection (*6*), distributed with ANTs 2.3.3 (*7*) (RRID:SCR_004757), and used as T1w-reference throughout the workflow. The T1w-reference was then skull-stripped with a Nipype implementation of the antsBrainExtraction.sh workflow (from ANTs), using OASIS30ANTs as target template. Brain tissue segmentation of cerebrospinal fluid (CSF), white-matter (WM) and gray-matter (GM) was performed on the brain-extracted T1w using fast (FSL 6.0.5.1:57b01774, RRID:SCR_002823) (*8*). Brain surfaces were reconstructed using recon-all (FreeSurfer 7.3.2, RRID:SCR_001847) (*9*), and the brain mask estimated previously was refined with a custom variation of the method to reconcile ANTs-derived and FreeSurfer-derived segmentations of the cortical gray-matter of Mindboggle (RRID:SCR_002438) (*10*).

Grayordinate “dscalar” files (*11*) containing 91k samples were also generated using the highest-resolution *fsaverage* as an intermediate standardized surface space. Volume-based spatial normalization to two standard spaces (MNI152NLin2009cAsym, MNI152NLin6Asym) was performed through nonlinear registration with *antsRegistration* (ANTs 2.3.3), using brain-extracted versions of both T1w reference and the T1w template. The following templates were selected for spatial normalization: ICBM 152 Nonlinear Asymmetrical template version 2009c (RRID:SCR_008796 (*12*); TemplateFlow ID: MNI152NLin2009cAsym), FSL’s MNI ICBM 152 non-linear 6th Generation Asymmetric Average Brain Stereotaxic Registration Model (RRID:SCR_002823 (*13*); TemplateFlow ID: MNI152NLin6Asym).

#### Functional data preprocessing

For each of the 10 BOLD runs per participant (5 encoding and 5 retrieval), the following preprocessing was performed. First, a reference volume and its skull-stripped version were generated using a custom methodology of fMRIPrep. Head-motion parameters with respect to the BOLD reference (transformation matrices, and six corresponding rotation and translation parameters) were estimated before spatiotemporal filtering using mcflirt (*14*) (FSL 6.0.5.1:57b01774). The estimated fieldmap was then aligned with rigid-registration to the target EPI (echo-planar imaging) reference run. The field coefficients were mapped on to the reference EPI using the transform. BOLD runs were slice-time corrected to 0.952s (0.5 of slice acquisition range 0s-1.9s) using 3dTshift from AFNI (*15*) (RRID:SCR_005927). The BOLD reference was then co-registered to the T1w reference using bbregister (FreeSurfer) which implements boundary-based registration (*16*). Co-registration was configured with six degrees of freedom.

Several confounds were calculated based on the preprocessed BOLD: framewise displacement (FD), DVARS and three region-wise global signals. FD was computed using two formulations, following Power (*17*) (absolute sum of relative motions) and Jenkinson (*14*) (relative root mean square displacement between affines). FD and DVARS were calculated for each functional run, using their implementations in Nipype (following the definitions by (*17*)). The three global signals were extracted within the CSF, WM, and whole-brain masks. Additionally, a set of physiological regressors were extracted to allow for component-based noise correction (*18*) (CompCor). Principal components were estimated after high-pass filtering the preprocessed BOLD time-series (using a discrete cosine filter with 128s cut-off) for the two CompCor variants: temporal (tCompCor) and anatomical (aCompCor). tCompCor components were then calculated from the top 2% variable voxels within the brain mask. For aCompCor, three probabilistic masks (CSF, WM and combined CSF+WM) were generated in anatomical space. The implementation differed from that of Behzadi et al. (*18*) in that instead of eroding the masks by two pixels on BOLD space, a mask of pixels that likely contain a volume fraction of GM was subtracted from the aCompCor masks. This mask was obtained by dilating a GM mask extracted from the FreeSurfer’s aseg segmentation, and it ensured components were not extracted from voxels containing a minimal fraction of GM. Finally, these masks were resampled into BOLD space and binarized by thresholding at 0.99 (as in the original implementation). Components were also calculated separately within the WM and CSF masks. For each CompCor decomposition, the k components with the largest singular values were retained, such that the retained components’ time-series were sufficient to explain 50 percent of variance across the nuisance mask (CSF, WM, combined, or temporal). The remaining components were dropped from consideration.

The head-motion estimates calculated in the correction step were also placed within the corresponding confounds file. The confound time-series derived from head motion estimates and global signals were expanded with the inclusion of temporal derivatives and quadratic terms for each (*19*). Frames that exceeded a threshold of 0.9 mm FD or 3.0 standardized DVARS were annotated as motion outliers. Additional nuisance time-series were calculated by means of principal components analysis of the signal found within a thin band (crown) of voxels around the edge of the brain, as proposed by (*20*). The BOLD time-series were resampled into standard space, generating a preprocessed BOLD run in MNI152NLin2009cAsym space. First, a reference volume and its skull-stripped version were generated using a custom methodology of fMRIPrep. The BOLD time-series were resampled onto the following surfaces (FreeSurfer reconstruction nomenclature): fsnative, fsaverage5, fsaverage. Grayordinates files (*11*) containing 91k samples were also generated using the highest-resolution fsaverage as intermediate standardized surface space. All resamplings can be performed with a single interpolation step by composing all the pertinent transformations (i.e. head-motion transform matrices, susceptibility distortion correction when available, and co-registrations to anatomical and output spaces). Gridded (volumetric) resamplings were performed using antsApplyTransforms (ANTs), configured with Lanczos interpolation to minimize the smoothing effects of other kernels (*21*). Non-gridded (surface) resamplings were performed using mri_vol2surf (FreeSurfer).

Many internal operations of fMRIPrep use Nilearn 0.9.1 (RRID:SCR_001362) (*22*), mostly within the functional processing workflow. For more details of the pipeline, see the section corresponding to workflows in fMRIPrep’s documentation.

## Supplementary tables

**Table S1.** Cortical brain regions exhibit significant category effects (n = 166; related to Fig. 1B).

| Contrast | Label index | Area (mm<sup>2</sup>) | DKT_overlap |
| --- | --- | --- | --- |
| **Face**<br>**>**<br>**Place** | 1 | 1421.36 | L_precuneus (74.1%); L_isthmuscingulate (25.9%); |
|  | 2 | 4393.79 | L_superiorfrontal (63.2%); L_medialorbitofrontal (13.1%);<br>L_rostralanteriorcingulate (10.6%); L_rostralmiddlefrontal (6.0%);<br>L_frontalpole (5.5%); |
|  | 3 | 11917.38 | L_superiortemporal (23.7%); L_middletemporal (15.9%);<br>L_inferiorparietal (13.3%); L_bankssts (7.3%); L_fusiform (6.4%);<br>L_lateraloccipital (6.2%); L_lateralorbitofrontal (5.5%); |
|  | 4 | 245.14 | L_lingual (100.0%); |
|  | 5 | 1315.14 | R_precuneus (76.1%); R_isthmuscingulate (23.9%); |
|  | 6 | 20183.86 | R_superiorfrontal (16.7%); R_middletemporal (13.2%);<br>R_inferiorparietal (10.6%); R_superiortemporal (10.1%);<br>R_fusiform (6.3%); R_medialorbitofrontal (6.0%);<br>R_lateraloccipital (6.0%); |
|  | 7 | 408.96 | R_lingual (100.0%); |
| **Face**<br>**<**<br>**Place** | 1 | 41115.71 | L_superiorparietal (10.9%); L_precentral (8.8%); L_lateraloccipital (8.0%);<br>L_superiorfrontal (7.8%); L_rostralmiddlefrontal (6.8%);<br>L_inferiorparietal (6.8%); L_precuneus (6.2%); L_inferiortemporal (5.4%);<br>L_supramarginal (5.1%); |
|  | 2 | 2969.92 | R_precentral (39.4%); R_parsopercularis (31.1%); R_insula (18.1%);<br>R_lateralorbitofrontal (5.2%); |
|  | 3 | 22558.92 | R_superiorparietal (19.8%); R_lateraloccipital (13.8%);<br>R_inferiorparietal (11.5%); R_precuneus (10.8%); R_lingual (10.1%);<br>R_supramarginal (9.6%); R_fusiform (7.2%); |
|  | 4 | 4296.19 | R_superiorfrontal (62.8%); R_caudalmiddlefrontal (17.2%);<br>R_precentral (11.9%); R_caudalanteriorcingulate (6.4%); |
|  | 5 | 1466.17 | R_inferiortemporal (82.4%); R_middletemporal (15.7%); |
|  | 6 | 2170.04 | R_rostralmiddlefrontal (90.5%); |
|  | 7 | 169.13 | R_superiortemporal (100.0%); |

Note: DKT = Desikan-Killiany-Tourville; This table was generated using ciftify\_statclust\_report (https://github.com/edickie/ciftify/blob/master/ciftify/bin/ciftify_statclust_report.py).

**Table S2.**
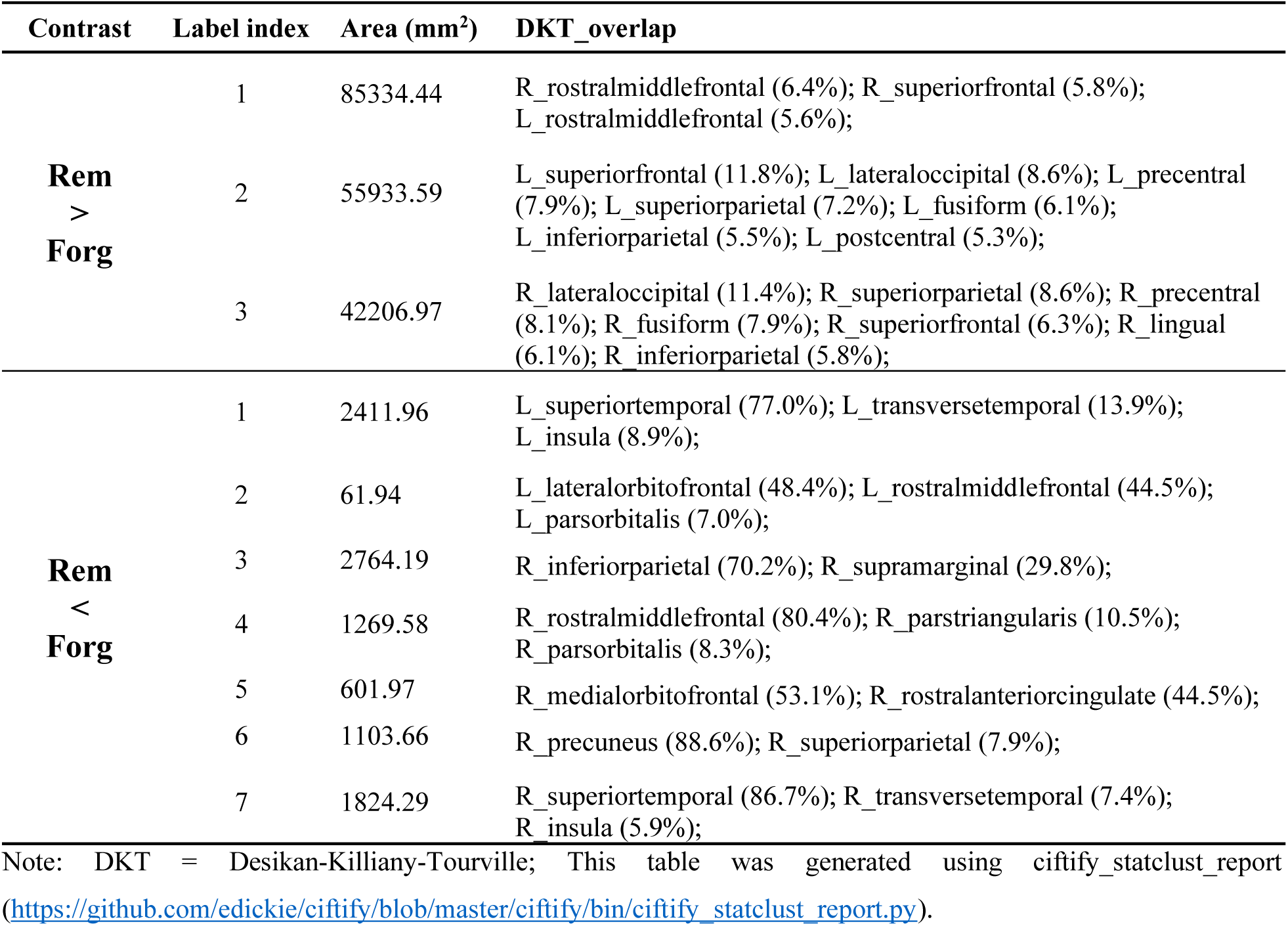
Cortical brain regions exhibit significant subsequent memory effects (n = 166; related to Fig. 1C).

| Contrast | Label index | Area (mm<sup>2</sup>) | DKT_overlap |
| --- | --- | --- | --- |
| **Rem**<br>**>**<br>**Forg** | 1 | 85334.44 | R_rostralmiddlefrontal (6.4%); R_superiorfrontal (5.8%);<br>L_rostralmiddlefrontal (5.6%); |
|  | 2 | 55933.59 | L_superiorfrontal (11.8%); L_lateraloccipital (8.6%); L_precentral<br>(7.9%); L_superiorparietal (7.2%); L_fusiform (6.1%);<br>L_inferiorparietal (5.5%); L_postcentral (5.3%); |
|  | 3 | 42206.97 | R_lateraloccipital (11.4%); R_superiorparietal (8.6%); R_precentral<br>(8.1%); R_fusiform (7.9%); R_superiorfrontal (6.3%); R_lingual<br>(6.1%); R_inferiorparietal (5.8%); |
| **Rem**<br>**<**<br>**Forg** | 1 | 2411.96 | L_superiortemporal (77.0%); L_transversetemporal (13.9%);<br>L_insula (8.9%); |
|  | 2 | 61.94 | L_lateralorbitofrontal (48.4%); L_rostralmiddlefrontal (44.5%);<br>L_parsorbitalis (7.0%); |
|  | 3 | 2764.19 | R_inferiorparietal (70.2%); R_supramarginal (29.8%); |
|  | 4 | 1269.58 | R_rostralmiddlefrontal (80.4%); R_parstriangularis (10.5%);<br>R_parsorbitalis (8.3%); |
|  | 5 | 601.97 | R_medialorbitofrontal (53.1%); R_rostralanteriorcingulate (44.5%); |
|  | 6 | 1103.66 | R_precuneus (88.6%); R_superiorparietal (7.9%); |
|  | 7 | 1824.29 | R_superiortemporal (86.7%); R_transversetemporal (7.4%);<br>R_insula (5.9%); |

Note: DKT = Desikan-Killiany-Tourville; This table was generated using ciftify\_statclust\_report (https://github.com/edickie/ciftify/blob/master/ciftify/bin/ciftify_statclust_report.py).

**Table S3.** Summary table of the LMM of age × memory (Rem vs Forg) × region (Face- vs Place-selective regions) predicting neural selectivity (n = 156; related to Fig. 2A).

| Neural selectivity ~ (age+educ+sex)*memory*region + (1\|subject) |  |  |  |  |
| --- | --- | --- | --- | --- |
| term | NumDF | DenDF | F | p |
| **age** | **1** | **152** | **7.355** | **0.007** |
| educ | 1 | 152 | 1.819 | 0.179 |
| sex | 1 | 152 | 7.938 | 0.005 |
| **memory** | **1** | **456** | **14.76** | **0** |
| **region** | **1** | **456** | **8.588** | **0.004** |
| **age:memory** | **1** | **456** | **10.25** | **0.001** |
| educ:memory | 1 | 456 | 0.56 | 0.455 |
| sex:memory | 1 | 456 | 8.608 | 0.004 |
| **age:region** | **1** | **456** | **6.756** | **0.01** |
| educ:region | 1 | 456 | 0.086 | 0.77 |
| sex:region | 1 | 456 | 0.15 | 0.699 |
| memory:region | 1 | 456 | 0.002 | 0.966 |
| age:memory:region | 1 | 456 | 0.323 | 0.57 |
| educ:memory:region | 1 | 456 | 1.219 | 0.27 |
| sex:memory:region | 1 | 456 | 0.148 | 0.701 |

Note: NumDF = numerator degree of freedom, DenDF = denominator degree of freedom, educ = education.

**Table S4.** Summary table of the LMM of age × category (Face vs Place) × region (Face-vs Place-selective regions) predicting preferred/non-preferred neural activity on remembered trials (n = 156; related to Fig. 2B).

| Neural activity ~ (age+educ+sex)*category*region + (1\|subject) |  |  |  |  |
| --- | --- | --- | --- | --- |
| term | NumDF | DenDF | F | p |
| **age** | **1** | **152** | **6.306** | **0.013** |
| sex | 1 | 152 | 11.146 | 0.001 |
| educ | 1 | 152 | 1.124 | 0.291 |
| category | 1 | 456 | 1.608 | 0.205 |
| region | 1 | 456 | 1.684 | 0.195 |
| age:category | 1 | 456 | 1.042 | 0.308 |
| sex:category | 1 | 456 | 0.075 | 0.785 |
| educ:category | 1 | 456 | 0.533 | 0.466 |
| age:region | 1 | 456 | 0.245 | 0.621 |
| sex:region | 1 | 456 | 23.177 | 0 |
| educ:region | 1 | 456 | 0.775 | 0.379 |
| **category:region** | **1** | **456** | **8.449** | **0.004** |
| **age:category:region** | **1** | **456** | **4.838** | **0.028** |
| sex:category:region | 1 | 456 | 4.653 | 0.032 |
| educ:category:region | 1 | 456 | 0.402 | 0.527 |

Note: NumDF = numerator degree of freedom, DenDF = denominator degree of freedom, educ = education.

**Table S5.** Summary table of the LMM of DAN SME × memory (Rem vs Forg) × region (Face-vs Place-selective regions) predicting neural selectivity (n = 156; related to Fig. 2C).

| Neural selectivity ~ sme_dan * memory * region + age + sex + educ + (1\|subject) |  |  |  |  |
| --- | --- | --- | --- | --- |
| term | NumDF | DenDF | F | p |
| **sme_dan** | **1** | **151** | **9.632** | **0.002** |
| memory | 1 | 462 | 46.826 | 0 |
| region | 1 | 462 | 41.504 | 0 |
| age | 1 | 151 | 5.911 | 0.016 |
| sex | 1 | 151 | 6.645 | 0.011 |
| educ | 1 | 151 | 1.174 | 0.28 |
| **sme_dan:memory** | **1** | **462** | **18.92** | **0** |
| sme_dan:region | 1 | 462 | 1.732 | 0.189 |
| **memory:region** | **1** | **462** | **4.165** | **0.042** |
| sme_dan:memory:region | 1 | 462 | 1.318 | 0.252 |

Note: NumDF = numerator degree of freedom, DenDF = denominator degree of freedom, educ = education, sme\_dan = subsequent memory effect in dorsal attention network.

**Table S6.** Summary table of the LMM of DAN SME × category (Face vs Place) × region (Face-vs Place-selective regions) predicting preferred/non-preferred neural activity on remembered trials (n = 156; related to Fig. 2D).

| <u>Neural activity ~ sme dan * category * region + age + sex + educ + (1\|subject)</u> |  |  |  |  |
| --- | --- | --- | --- | --- |
| **term** | **NumDF** | **DenDF** | **F** | **p** |
| sme_dan | 1 | 151 | 72.652 | 0 |
| category | 1 | 462 | 0.162 | 0.688 |
| region | 1 | 462 | 211.289 | 0 |
| age | 1 | 151 | 4.386 | 0.038 |
| sex | 1 | 151 | 10.2 | 0.002 |
| educ | 1 | 151 | 0.216 | 0.643 |
| sme_dan:category | 1 | 462 | 0.722 | 0.396 |
| sme_dan:region | 1 | 462 | 14.059 | 0 |
| category:region | 1 | 462 | 245.168 | 0 |
| **sme_dan:category:region** | **1** | **462** | **6.618** | **0.01** |
| **Region = Face-selective ROIs** |  |  |  |  |
| sme_dan | 1 | 222.05 | 86.651 | 0 |
| category | 1 | 462 | 230.058 | 0 |
| age | 1 | 151 | 4.386 | 0.038 |
| sex | 1 | 151 | 10.2 | 0.002 |
| educ | 1 | 151 | 0.216 | 0.643 |
| sme_dan:category | 1 | 462 | 1.484 | 0.224 |
| **Region = Place-selective ROIs** |  |  |  |  |
| sme_dan | 1 | 222.05 | 37.332 | 0 |
| category | 1 | 462 | 236.072 | 0 |
| age | 1 | 151 | 4.386 | 0.038 |
| sex | 1 | 151 | 10.2 | 0.002 |
| educ | 1 | 151 | 0.216 | 0.643 |
| **sme_dan:category** | **1** | **462** | **5.856** | **0.016** |

Note: NumDF = numerator degree of freedom, DenDF = denominator degree of freedom, educ = education, sme\_dan = subsequent memory effect in dorsal attention network.

**Table S7.** Summary table of the LMM of plasma assay (log-transformed pTau_181_ and Aβ_42_/Aβ_40_) × memory (Rem vs Forg) × region (Face-vs Place-selective regions) predicting neural selectivity (related to Fig. 2E).

| **Plasma biomarkers** | **term** | **NumDF** | **DenDF** | **F** | **p** |
| --- | --- | --- | --- | --- | --- |
| *Neural selectivity ~ pTau181\*memory\*region + age + sex + educ + (1/subject)* |  |  |  |  |  |
| pTau181<br>(log)<br>n = 137 | pTau181 | 1 | 132 | 0.266 | 0.607 |
|  | memory | 1 | 405 | 69.426 | 0 |
|  | region | 1 | 405 | 72.832 | 0 |
|  | age | 1 | 132 | 5.937 | 0.016 |
|  | sex | 1 | 132 | 5.406 | 0.022 |
|  | educ | 1 | 132 | 0.809 | 0.37 |
|  | pTau181:memory | 1 | 405 | 3.561 | 0.06 |
|  | **pTau181:region** | **1** | **405** | **14.797** | **0** |
|  | memory:region | 1 | 405 | 0.511 | 0.475 |
|  | pTau181:memory:region | 1 | 405 | 0.037 | 0.847 |
| *Neural selectivity ~ AB42/AB40\*memory\*region + age + sex + educ + (1/subject)* |  |  |  |  |  |
| AB42/AB40<br>n = 136 | AB42/AB40 | 1 | 131 | 3.038 | 0.084 |
|  | memory | 1 | 402 | 0.002 | 0.968 |
|  | region | 1 | 402 | 0.06 | 0.807 |
|  | age | 1 | 131 | 6.531 | 0.012 |
|  | sex | 1 | 131 | 4.639 | 0.033 |
|  | educ | 1 | 131 | 0.379 | 0.539 |
|  | AB42/AB40:memory | 1 | 402 | 2.377 | 0.124 |
|  | AB42/AB40:region | 1 | 402 | 2.311 | 0.129 |
|  | memory:region | 1 | 402 | 0.317 | 0.574 |
|  | AB42/AB40:memory:region | 1 | 402 | 0.545 | 0.461 |

Note: NumDF = numerator degree of freedom, DenDF = denominator degree of freedom, educ = education.

**Table S8.** Summary table of the LMM of plasma pTau_181_ × category (Face vs Place) predicting preferred/non-preferred neural activity on remembered trials in place-selective regions (n = 137; related to Fig. 2F).

|  |
| --- |
| *Neural activity ~ pTau181 \* category + age + sex + educ + (1\|subj_id)* |

| term | NumDF | DenDF | F | p |
| --- | --- | --- | --- | --- |
| pTau181 | 1 | 132 | 0.175 | 0.676 |
| category | 1 | 135 | 260.506 | 0 |
| age | 1 | 132 | 8.416 | 0.004 |
| sex | 1 | 132 | 2.132 | 0.147 |
| educ | 1 | 132 | 0.194 | 0.66 |
| **pTau181:category** | **1** | **135** | **10.392** | **0.002** |

Note: NumDF = numerator degree of freedom, DenDF = denominator degree of freedom, educ = education.

**Table S9.** Summary table of the LMM of CSF assay (pTau_181_ and Aβ_42_/Aβ_40_) × memory (Rem vs Forg) × region (Face-vs Place-selective regions) predicting neural selectivity (n = 115; related to Fig. S6).

| CSF biomarkers | term | NumDF | DenDF | F | p |
| --- | --- | --- | --- | --- | --- |
| *Neural selectivity ~ pTau181\*memory\*region + age + sex + educ + (1/subject)* |  |  |  |  |  |
| pTau181 (log) | pTau181 | 1 | 110 | 0.263 | 0.609 |
|  | memory | 1 | 339 | 4.599 | 0.033 |
|  | region | 1 | 339 | 9.862 | 0.002 |
|  | age | 1 | 110 | 5.075 | 0.026 |
|  | sex | 1 | 110 | 5.775 | 0.018 |
|  | educ | 1 | 110 | 1.274 | 0.261 |
|  | pTau181:memory | 1 | 339 | 0.771 | 0.381 |
|  | **pTau181:region** | **1** | **339** | **4.657** | **0.032** |
|  | memory:region | 1 | 339 | 0.003 | 0.954 |
|  | pTau181:memory:region | 1 | 339 | 0.066 | 0.797 |
| *Neural selectivity ~ AB42/AB40\*memory\*region + age + sex + educ + (1/subject)* |  |  |  |  |  |
| AB42/AB40 | AB42/AB40 | 1 | 110 | 0.052 | 0.821 |
|  | memory | 1 | 339 | 5.003 | 0.026 |
|  | region | 1 | 339 | 0.012 | 0.914 |
|  | age | 1 | 110 | 4.881 | 0.029 |
|  | sex | 1 | 110 | 6.077 | 0.015 |
|  | educ | 1 | 110 | 1.283 | 0.26 |
|  | AB42/AB40:memory | 1 | 339 | 0.208 | 0.649 |
|  | **AB42/AB40:region** | **1** | **339** | **4.228** | **0.041** |
|  | memory:region | 1 | 339 | 0.576 | 0.448 |
|  | AB42/AB40:memory:region | 1 | 339 | 0.125 | 0.723 |

Note: NumDF = numerator degree of freedom, DenDF = denominator degree of freedom, educ = education.

**Table S10.** Summary of hierarchical linear regression models predicting neural selectivity (n = 137; related to Figs. 3B & C).

|  | Formula | Excluded regressors | R<sup>2</sup> |
| --- | --- | --- | --- |
| Full model | NS ~ DAN + Age + AD + Sex + Educ | -- | 0.259*** |
| 1a | NS ~ Age + AD + Sex + Educ | DAN | 0.165*** |
| 1b | NS ~ DAN + AD + Sex + Educ | Age | 0.225*** |
| 1c | NS ~ DAN + Age + Sex + Educ | AD | 0.239*** |
| 2a | NS ~ DAN + Sex + Educ | Age & AD | 0.168*** |
| 2b | NS ~ AD + Sex + Educ | Age & DAN | 0.114** |
| 2c | NS ~ Age + Sex + Educ | DAN & AD | 0.144*** |

Note: NS = neural selectivity, DAN = DAN SME (subsequent memory effect in dorsal attention network), AD = plasma pTau<sub>181</sub>, Educ = years of education; \*\* p < .01, \*\*\* p < .001.

**Table S11.** Summary of hierarchical linear regression models predicting memory performance (n = 137; related to Figs. 5B & C).

|  | Formula | Excluded regressors | R<sup>2</sup> |
| --- | --- | --- | --- |
| Full model | Memory ~ NS + DAN + Age + AD + Sex + Educ | -- | 0.284*** |
| 1a | Memory ~ DAN + Age + AD + Sex + Educ | NS | 0.240*** |
| 1b | Memory ~ NS + Age + AD + Sex + Educ | DAN | 0.260*** |
| 1c | Memory ~ NS + DAN + AD + Sex + Educ | Age | 0.256*** |
| 1d | Memory ~ NS + DAN + Age + Sex + Educ | AD | 0.271*** |
| 2a | Memory ~ DAN + NS + Sex + Educ | Age & AD | 0.222*** |
| 3a | Memory ~ AD | NS, DAN, Age, Sex & Educ | 0.085*** |

Note: Memory = place associative $d'$ , NS = neural selectivity, DAN = DAN SME (subsequent memory effect in dorsal attention network), AD = plasma pTau<sub>181</sub>, Educ = years of education; \*\*\* p < .001.

**Table S12.**
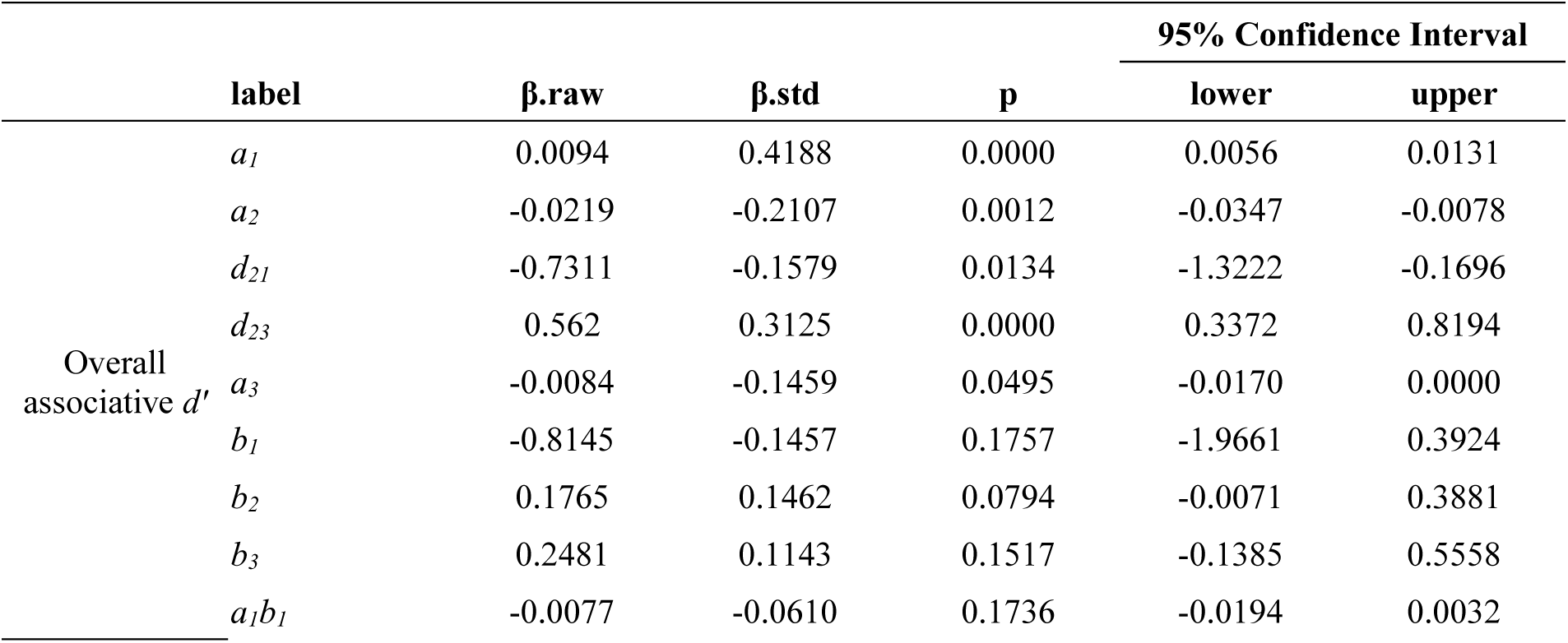
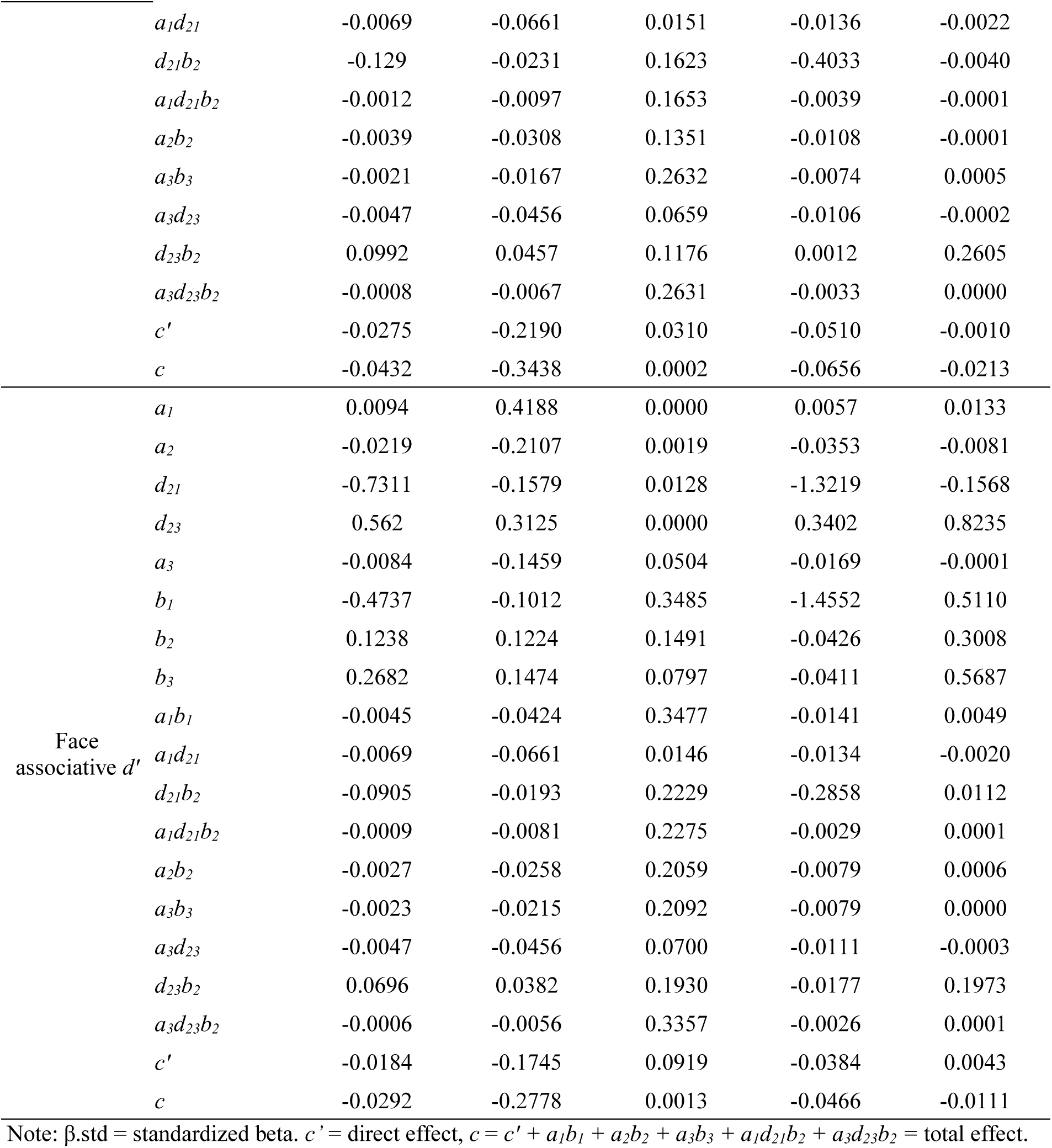
Results from SEM for overall associative *d’* and face associative *d’* (n = 137).

| | label | $\beta$.raw | $\beta$.std | p | 95% Confidence Interval | |
| --- | --- | --- | --- | --- | --- | --- |
|  |  |  |  |  | lower | upper |
| Overall<br>associative $d'$ | $a_1$ | 0.0094 | 0.4188 | 0.0000 | 0.0056 | 0.0131 |
| | $a_2$ | -0.0219 | -0.2107 | 0.0012 | -0.0347 | -0.0078 |
| | $d_{21}$ | -0.7311 | -0.1579 | 0.0134 | -1.3222 | -0.1696 |
| | $d_{23}$ | 0.562 | 0.3125 | 0.0000 | 0.3372 | 0.8194 |
| | $a_3$ | -0.0084 | -0.1459 | 0.0495 | -0.0170 | 0.0000 |
| | $b_1$ | -0.8145 | -0.1457 | 0.1757 | -1.9661 | 0.3924 |
| | $b_2$ | 0.1765 | 0.1462 | 0.0794 | -0.0071 | 0.3881 |
| | $b_3$ | 0.2481 | 0.1143 | 0.1517 | -0.1385 | 0.5558 |
| | $a_1b_1$ | -0.0077 | -0.0610 | 0.1736 | -0.0194 | 0.0032 |
| | $a_1d_{21}$ | -0.0069 | -0.0661 | 0.0151 | -0.0136 | -0.0022 |
| | $d_{21}b_2$ | -0.129 | -0.0231 | 0.1623 | -0.4033 | -0.0040 |
| | $a_1d_{21}b_2$ | -0.0012 | -0.0097 | 0.1653 | -0.0039 | -0.0001 |
| | $a_2b_2$ | -0.0039 | -0.0308 | 0.1351 | -0.0108 | -0.0001 |
| | $a_3b_3$ | -0.0021 | -0.0167 | 0.2632 | -0.0074 | 0.0005 |
| | $a_3d_{23}$ | -0.0047 | -0.0456 | 0.0659 | -0.0106 | -0.0002 |
| | $d_{23}b_2$ | 0.0992 | 0.0457 | 0.1176 | 0.0012 | 0.2605 |
| | $a_3d_{23}b_2$ | -0.0008 | -0.0067 | 0.2631 | -0.0033 | 0.0000 |
| | $c'$ | -0.0275 | -0.2190 | 0.0310 | -0.0510 | -0.0010 |
| | $c$ | -0.0432 | -0.3438 | 0.0002 | -0.0656 | -0.0213 |
| Face<br>associative $d'$ | $a_1$ | 0.0094 | 0.4188 | 0.0000 | 0.0057 | 0.0133 |
| | $a_2$ | -0.0219 | -0.2107 | 0.0019 | -0.0353 | -0.0081 |
| | $d_{21}$ | -0.7311 | -0.1579 | 0.0128 | -1.3219 | -0.1568 |
| | $d_{23}$ | 0.562 | 0.3125 | 0.0000 | 0.3402 | 0.8235 |
| | $a_3$ | -0.0084 | -0.1459 | 0.0504 | -0.0169 | -0.0001 |
| | $b_1$ | -0.4737 | -0.1012 | 0.3485 | -1.4552 | 0.5110 |
| | $b_2$ | 0.1238 | 0.1224 | 0.1491 | -0.0426 | 0.3008 |
| | $b_3$ | 0.2682 | 0.1474 | 0.0797 | -0.0411 | 0.5687 |
| | $a_1b_1$ | -0.0045 | -0.0424 | 0.3477 | -0.0141 | 0.0049 |
| | $a_1d_{21}$ | -0.0069 | -0.0661 | 0.0146 | -0.0134 | -0.0020 |
| | $d_{21}b_2$ | -0.0905 | -0.0193 | 0.2229 | -0.2858 | 0.0112 |
| | $a_1d_{21}b_2$ | -0.0009 | -0.0081 | 0.2275 | -0.0029 | 0.0001 |
| | $a_2b_2$ | -0.0027 | -0.0258 | 0.2059 | -0.0079 | 0.0006 |
| | $a_3b_3$ | -0.0023 | -0.0215 | 0.2092 | -0.0079 | 0.0000 |
| | $a_3d_{23}$ | -0.0047 | -0.0456 | 0.0700 | -0.0111 | -0.0003 |
| | $d_{23}b_2$ | 0.0696 | 0.0382 | 0.1930 | -0.0177 | 0.1973 |
| | $a_3d_{23}b_2$ | -0.0006 | -0.0056 | 0.3357 | -0.0026 | 0.0001 |
| | $c'$ | -0.0184 | -0.1745 | 0.0919 | -0.0384 | 0.0043 |
| | $c$ | -0.0292 | -0.2778 | 0.0013 | -0.0466 | -0.0111 |

Note: $\beta$ .std = standardized beta. $c'$ = direct effect, $c = c' + a_1b_1 + a_2b_2 + a_3b_3 + a_1d_{21}b_2 + a_3d_{23}b_2$ = total effect.

**Table S13.** Mediation results of age → neural selectivity → memory performance (overall, place, and face associative *d’*) (n = 156).

| | label | $\beta$.raw | $\beta$.std | pvalue | 95% Confidence Interval | |
| --- | --- | --- | --- | --- | --- | --- |
|  |  |  |  |  | lower | upper |
| Place<br>associative $d'$ | $b$ | 0.366 | 0.308 | <.001 | 0.202 | 0.551 |
| | $c'$ | -0.028 | -0.229 | 0.009 | -0.05 | -0.008 |
|  | *a* | -0.029 | -0.276 | <.001 | -0.041 | -0.015 |
|  | ***ab*** | **-0.011** | **-0.085** | **0.003** | **-0.019** | **-0.005** |
|  | *c'+ab* | -0.039 | -0.314 | <.001 | -0.061 | -0.017 |
| Face<br>associative *d'* | *b* | 0.233 | 0.22 | 0.005 | 0.08 | 0.409 |
|  | *c'* | -0.02 | -0.178 | 0.033 | -0.038 | -0.001 |
|  | *a* | -0.029 | -0.276 | <.001 | -0.041 | -0.015 |
|  | ***ab*** | **-0.007** | **-0.061** | **0.024** | **-0.014** | **-0.002** |
|  | *c'+ab* | -0.026 | -0.239 | 0.003 | -0.043 | -0.009 |
| Overall<br>associative *d'* | *b* | 0.259 | 0.21 | 0.006 | 0.08 | 0.444 |
|  | *c'* | -0.033 | -0.253 | 0.003 | -0.053 | -0.011 |
|  | *a* | -0.029 | -0.276 | <.001 | -0.042 | -0.016 |
|  | ***ab*** | **-0.007** | **-0.058** | **0.023** | **-0.015** | **-0.002** |
|  | *c'+ab* | -0.04 | -0.311 | <.001 | -0.061 | -0.018 |

Note: $\beta$ .std = standardized beta. *c'* = direct effect, *ab* = indirect effect, *c'+ab* = total effect.

**Table S14.**
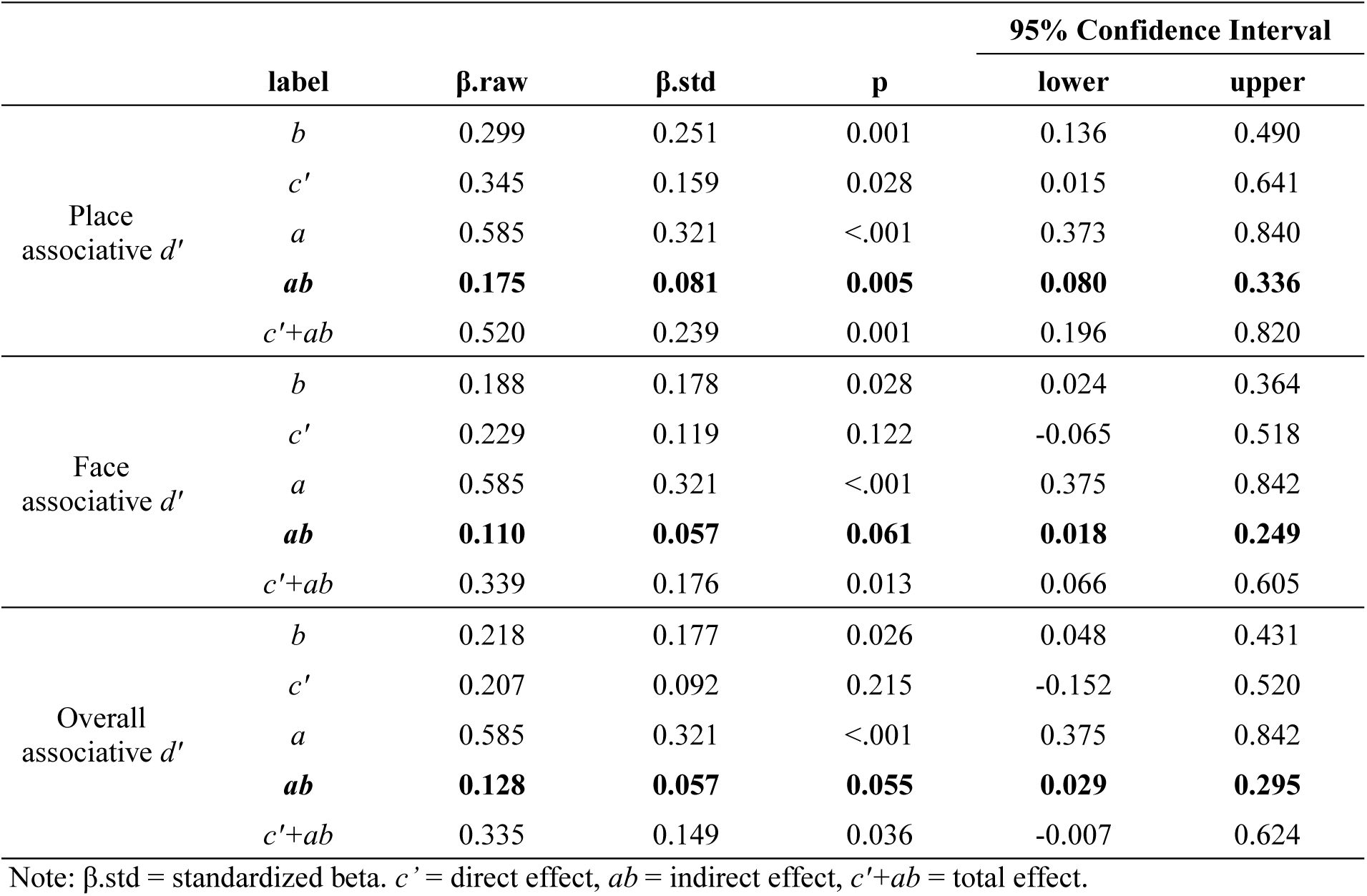
Mediation results of DAN SME → neural selectivity → memory performance (overall, place, and face associative *d’*) (n = 156).

**Table S15.** Results from SEM for delayed recall composite score (n = 137).

| | label | $\beta$.raw | $\beta$.std | p | 95% Confidence Interval | |
| --- | --- | --- | --- | --- | --- | --- |
|  |  |  |  |  | lower | upper |
| Delayed recall composite score (Z) | $a_1$ | 0.0094 | 0.4188 | 0.0000 | 0.0057 | 0.0132 |
| | $a_2$ | -0.0219 | -0.2107 | 0.0019 | -0.0358 | -0.0078 |
| | $d_{21}$ | -0.7311 | -0.1579 | 0.0148 | -1.3118 | -0.1199 |
| | $d_{23}$ | 0.5620 | 0.3125 | 0.0000 | 0.3429 | 0.8252 |
| | $a_3$ | -0.0084 | -0.1459 | 0.0495 | -0.0170 | 0.0001 |
| | $b_1$ | -0.1847 | -0.0314 | 0.7553 | -1.4875 | 0.8491 |
| | $b_2$ | 0.2104 | 0.1653 | 0.0330 | 0.0113 | 0.3962 |
| | $b_3$ | 0.3503 | 0.1530 | 0.0477 | -0.0075 | 0.6874 |
| | $a_1b_1$ | -0.0017 | -0.0131 | 0.7594 | -0.0150 | 0.0085 |
| | $a_1d_{21}$ | -0.0069 | -0.0661 | 0.0168 | -0.0135 | -0.0019 |
| | $d_{21}b_2$ | -0.1538 | -0.0261 | 0.1273 | -0.4280 | -0.0089 |
| | $a_1d_{21}b_2$ | -0.0015 | -0.0109 | 0.1246 | -0.0042 | -0.0001 |
| | $a_2b_2$ | -0.0046 | -0.0348 | 0.0803 | -0.0113 | -0.0006 |
| | $a_3b_3$ | -0.0030 | -0.0223 | 0.1842 | -0.0094 | 0.0000 |
| | $a_3d_{23}$ | -0.0047 | -0.0456 | 0.0634 | -0.0104 | -0.0002 |
| | $d_{23}b_2$ | 0.1183 | 0.0517 | 0.0624 | 0.0106 | 0.2657 |
| | $a_3d_{23}b_2$ | -0.0010 | -0.0075 | 0.1852 | -0.0033 | 0.0000 |
| | $c'$ | -0.0367 | -0.2772 | 0.0030 | -0.0590 | -0.0106 |
| | $c$ | -0.0485 | -0.3660 | 0.0000 | -0.0700 | -0.0250 |

Note: $\beta$ .std = standardized beta. $c'$ = direct effect, $c = c' + a_1b_1 + a_2b_2 + a_3b_3 + a_1d_{21}b_2 + a_3d_{23}b_2$ = total effect.

**Table S16.**
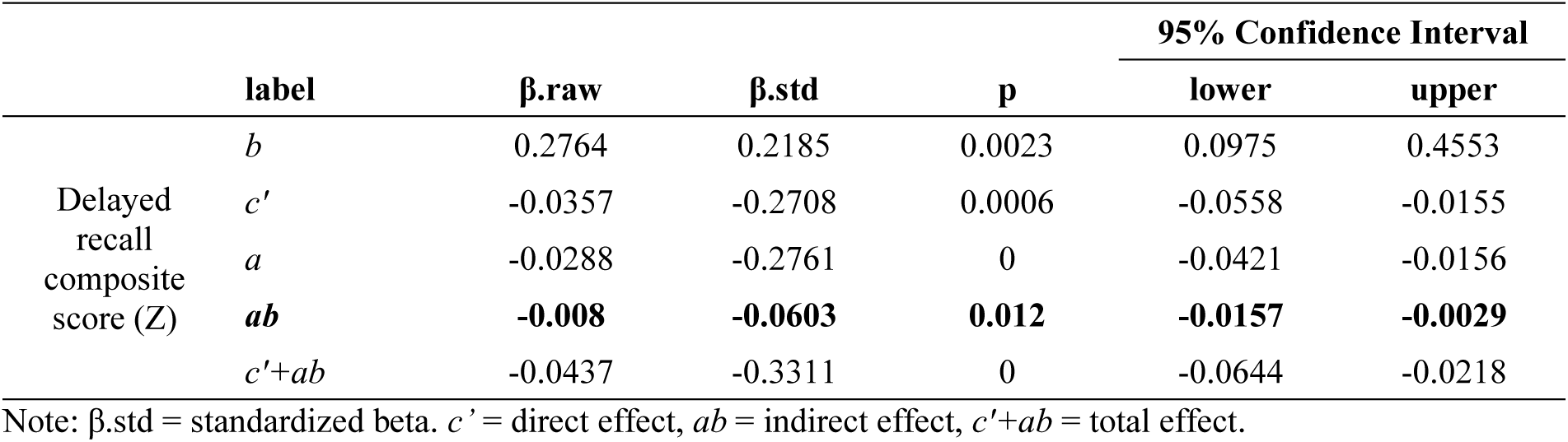
Mediation results of age → neural selectivity → delayed recall composite score (n = 156).

| | label | $\beta$.raw | $\beta$.std | p | 95% Confidence Interval | |
| --- | --- | --- | --- | --- | --- | --- |
|  |  |  |  |  | lower | upper |
| Delayed recall composite score (Z) | $b$ | 0.2764 | 0.2185 | 0.0023 | 0.0975 | 0.4553 |
| | $c'$ | -0.0357 | -0.2708 | 0.0006 | -0.0558 | -0.0155 |
| | $a$ | -0.0288 | -0.2761 | 0 | -0.0421 | -0.0156 |
|  | **$ab$** | **-0.008** | **-0.0603** | **0.012** | **-0.0157** | **-0.0029** |
| | $c'+ab$ | -0.0437 | -0.3311 | 0 | -0.0644 | -0.0218 |

Note: $\beta$ .std = standardized beta. $c'$ = direct effect, $ab$ = indirect effect, $c'+ab$ = total effect.

**Table S17.** Mediation results of top-down attention → neural selectivity → delayed recall composite score (n = 156).

|  |  |  |  |  | 95% Confidence Interval |  |
| --- | --- | --- | --- | --- | --- | --- |
| | label | $\beta$.raw | $\beta$.std | p | lower | upper |
| Delayed recall composite score (Z) | *b* | 0.215 | 0.17 | 0.024 | 0.027 | 0.403 |
|  | *c'* | 0.315 | 0.136 | 0.075 | -0.035 | 0.651 |
|  | *a* | 0.585 | 0.321 | 0 | 0.371 | 0.831 |
|  | ***ab*** | **0.126** | **0.055** | **0.049** | **0.022** | **0.28** |
|  | *c'+ab* | 0.441 | 0.191 | 0.008 | 0.105 | 0.757 |

Note: $\beta$ .std = standardized beta. *c'* = direct effect, *ab* = indirect effect, *c'+ab* = total effect.

**Table S18.**
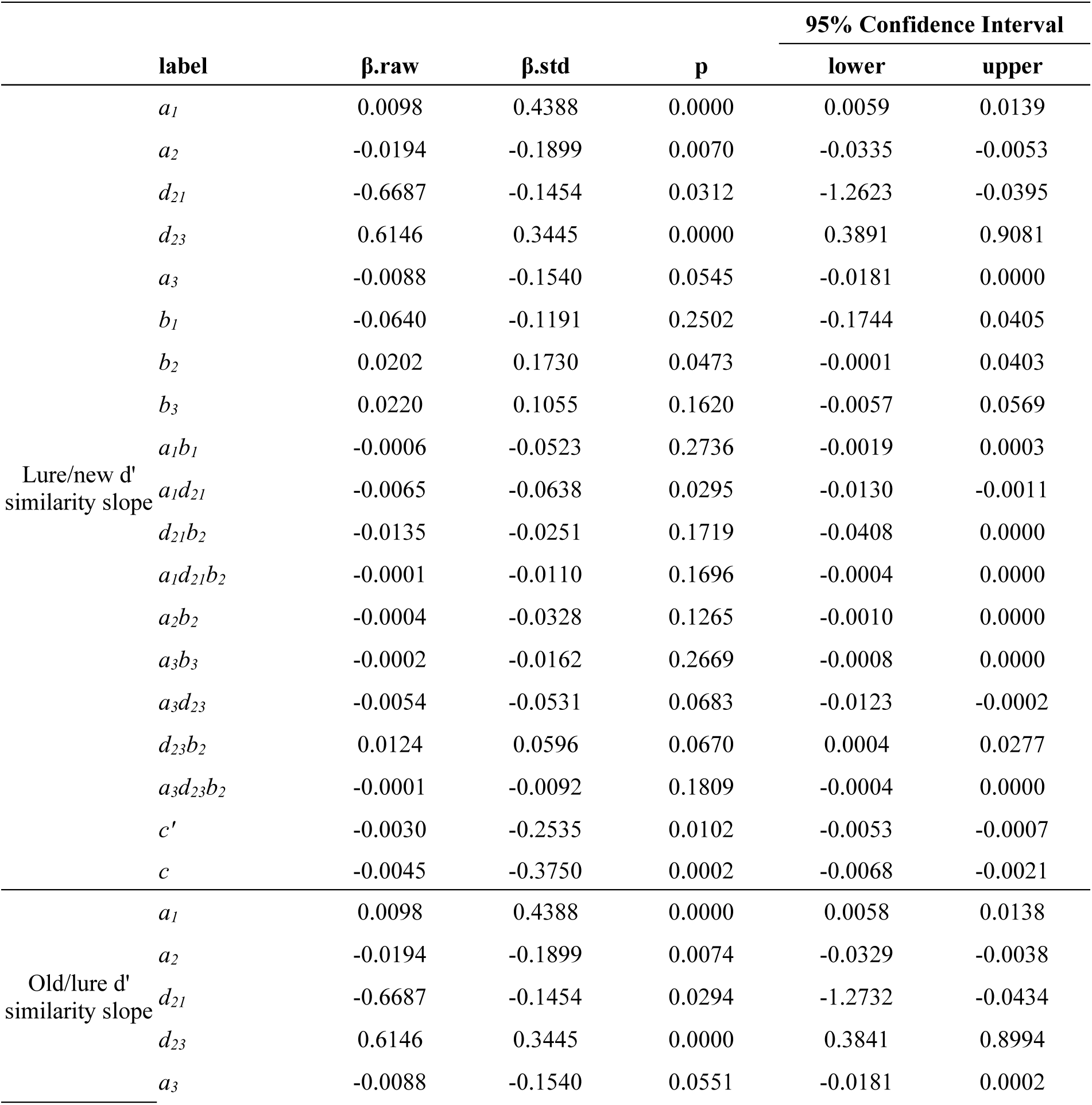
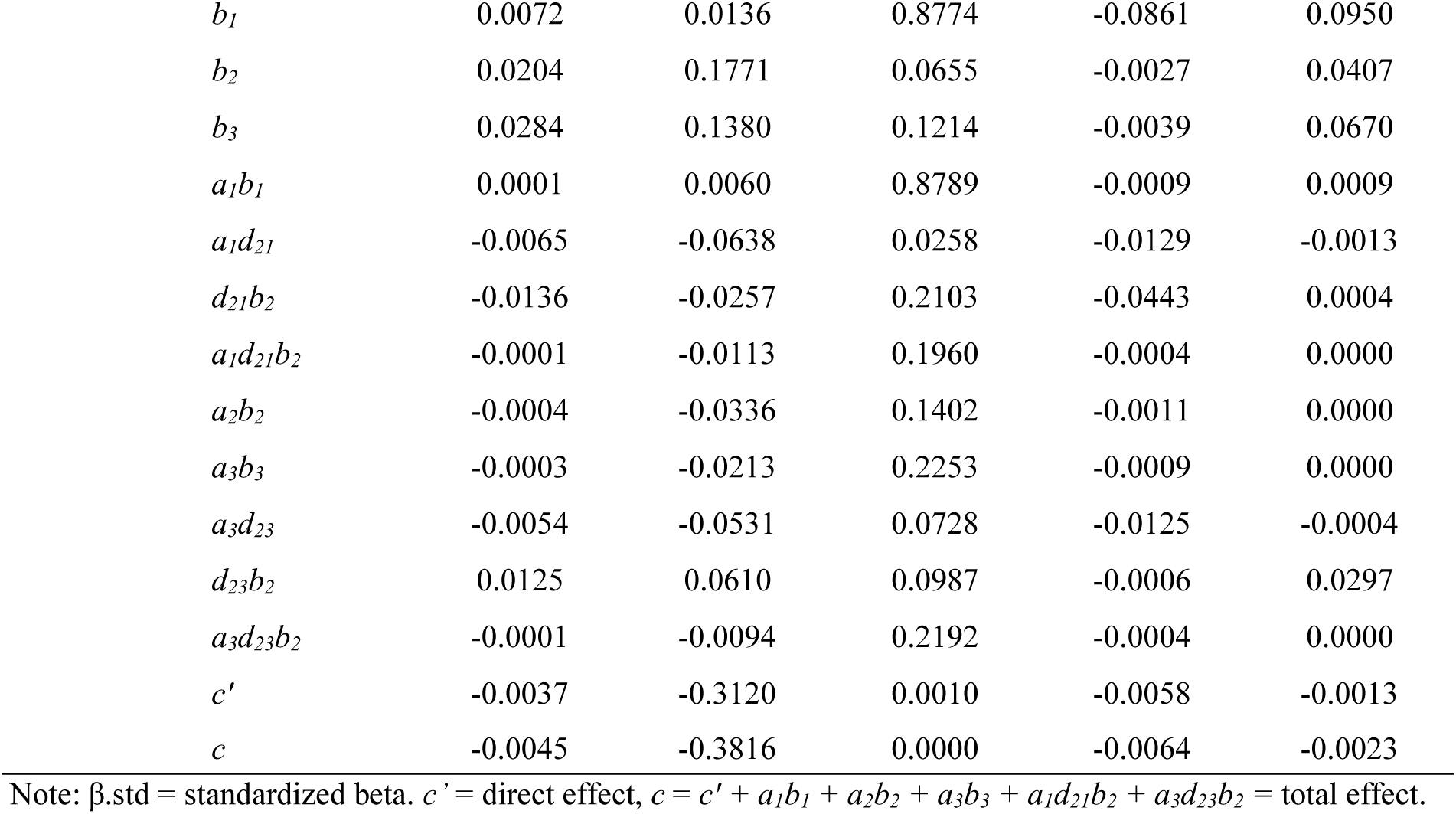
Results from SEM for mnemonic similarity task performance (n = 119).

**Table S19.** Mediation results of age → neural selectivity → mnemonic similarity task performance (n = 135).

| | label | $\beta$.raw | $\beta$.std | p | 95% Confidence Interval | |
| --- | --- | --- | --- | --- | --- | --- |
|  |  |  |  |  | lower | upper |
| Lure/new $d'$<br>similarity<br>slope | $b$ | 0.026 | 0.221 | 0.0039 | 0.0083 | 0.0439 |
| | $c'$ | -0.0034 | -0.2845 | 0.0036 | -0.0057 | -0.0011 |
| | $a$ | -0.0282 | -0.2746 | 0.0001 | -0.0416 | -0.0141 |
|  | **$ab$** | **-0.0007** | **-0.0607** | **0.0187** | **-0.0015** | **-0.0002** |
| | $c'+ab$ | -0.0042 | -0.3452 | 0.0003 | -0.0064 | -0.0018 |
| Old/lure $d'$<br>similarity<br>slope | $b$ | 0.0285 | 0.246 | 0.0024 | 0.0093 | 0.0467 |
| | $c'$ | -0.0032 | -0.2674 | 0.0014 | -0.0051 | -0.0012 |
| | $a$ | -0.0282 | -0.2746 | 0 | -0.0415 | -0.0144 |
|  | **$ab$** | **-0.0008** | **-0.0676** | **0.0165** | **-0.0016** | **-0.0002** |
| | $c'+ab$ | -0.004 | -0.335 | 0 | -0.0058 | -0.002 |

Note: $\beta$ .std = standardized beta. $c'$ = direct effect, $ab$ = indirect effect, $c'+ab$ = total effect.

**Table S20.**
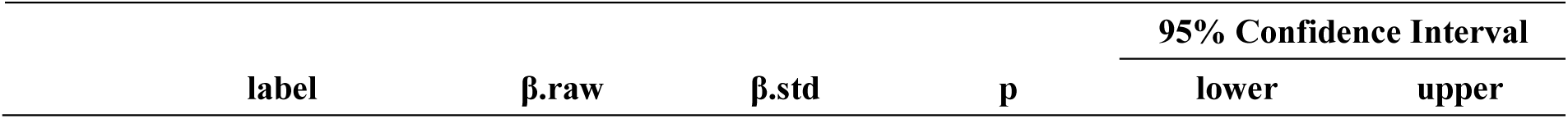
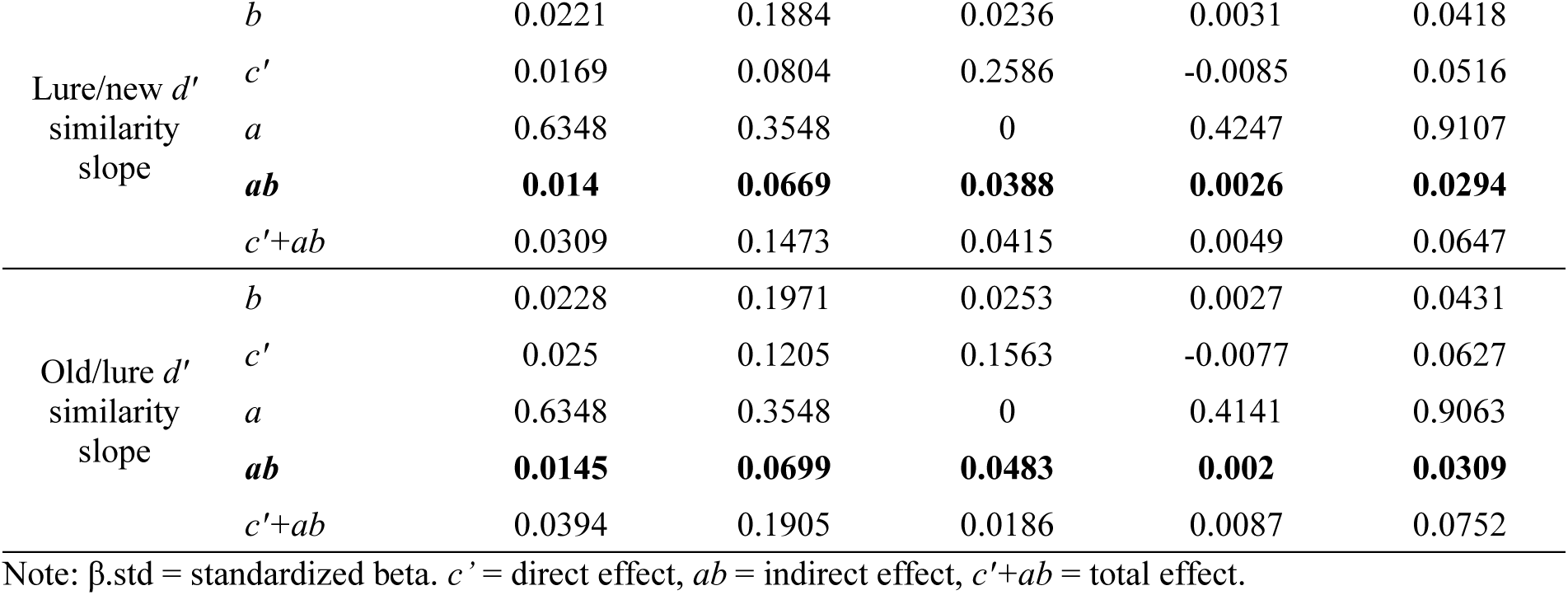
Mediation results of top-down attention → neural selectivity → mnemonic similarity task performance (n = 135).

## Supplementary Figures

**Fig. S1.**
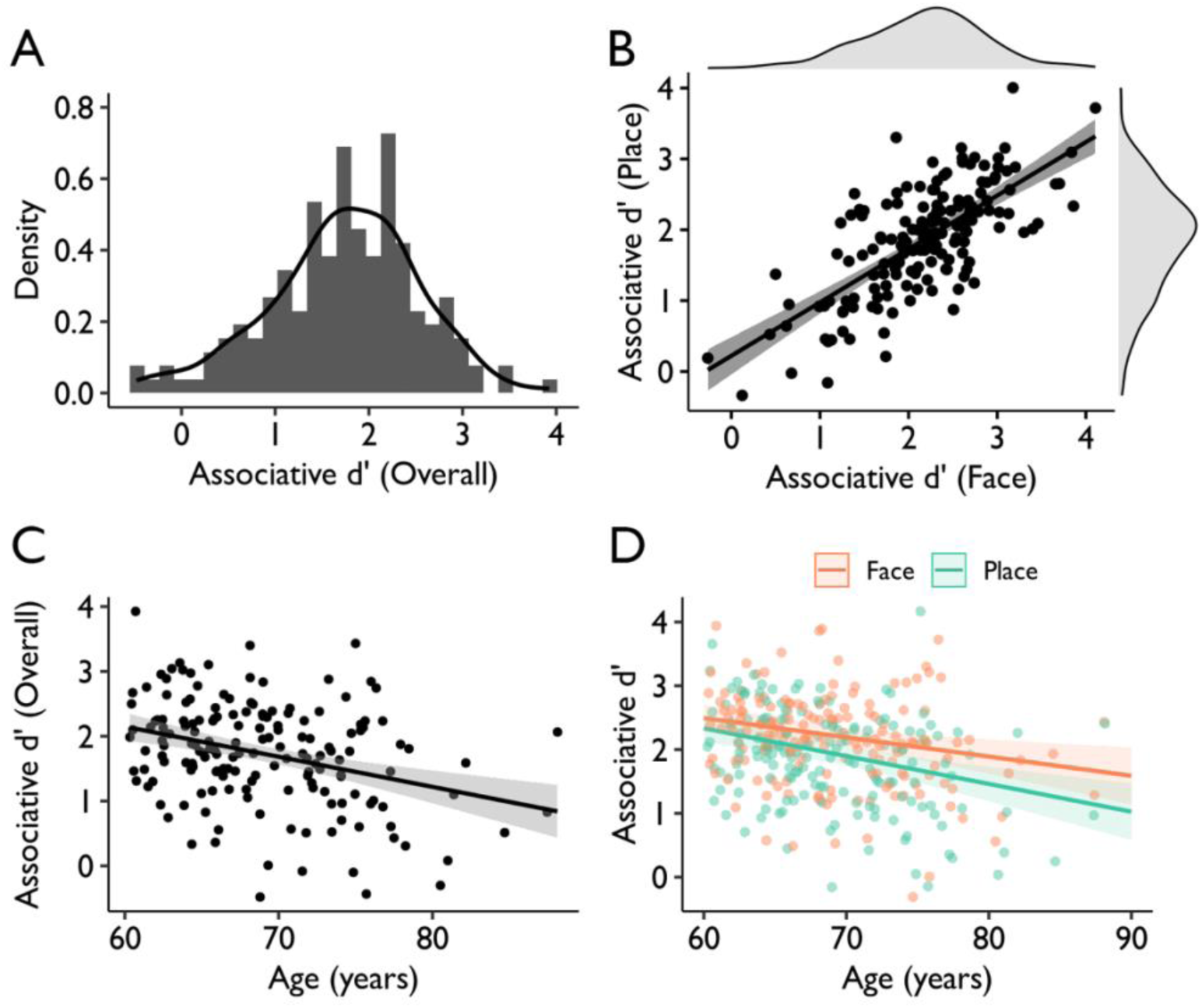
Behavioral results. (A) Distribution of overall associative *d’* performance. (B) Face and place associative memory were highly correlated (R = 0.71, p < 0.001). Overall (skew = -.33, kurtosis = .16), face (skew = -.29, kurtosis = .58), and place (skew = -.26, kurtosis = .04) associative *d’* followed a normal-like distribution. (C) Overall and (D) face (warm color) and place (cool color) memory performance declined with age.

**Fig. S2.**
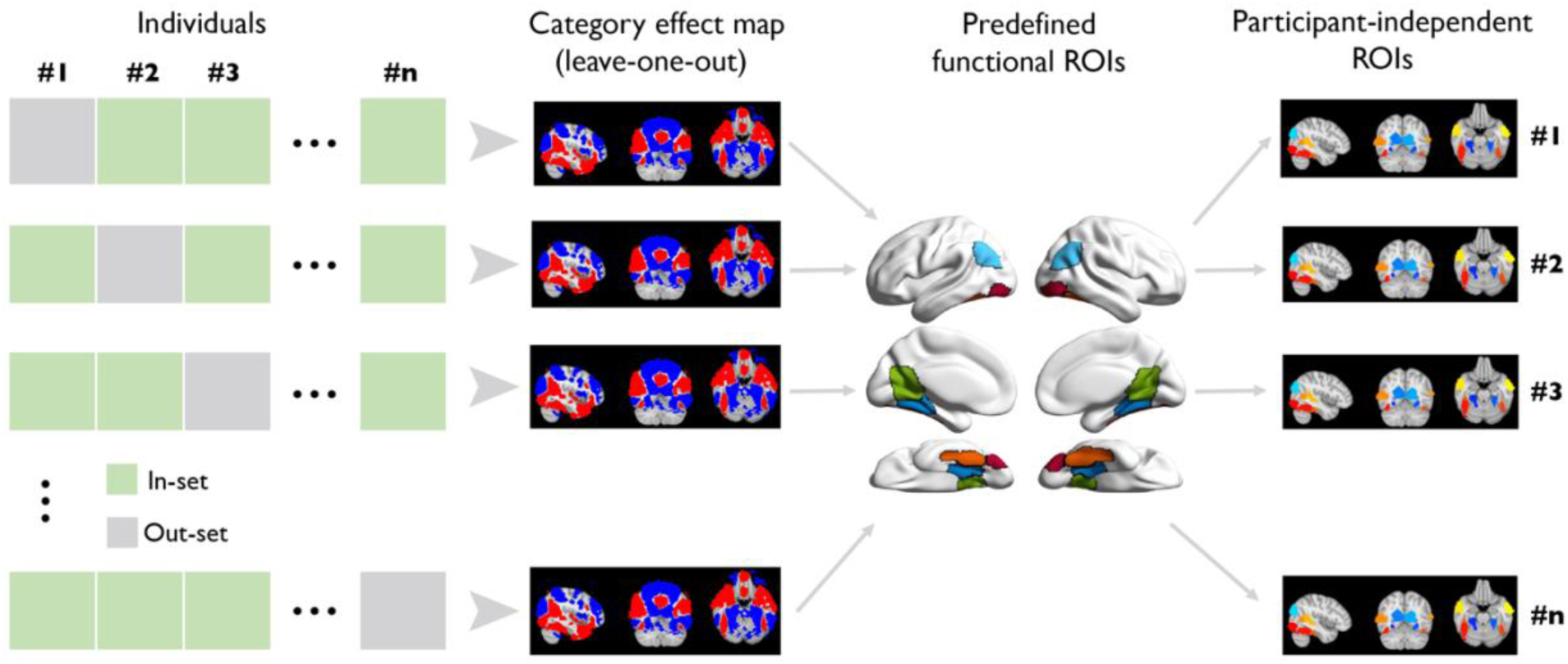
Leave-one-participant-out ROI delineation. For a given participant, we first performed a group-level analysis without the participant to obtain a whole-brain category effect in standard MNI space. A significant category effect map was extracted after controlling for family-wise error (FWE) with p < 0.05 using threshold-free cluster enhancement (TFCE) (Smith and Nichols, NI, 2009). Next, the resulting whole-brain map was intersected with the predefined functional ROIs, thus yielding study-specific and participant-independent ROIs. Finally, we transformed the study-specific ROIs to the given participant’s brain in native space.

**Fig. S3.**
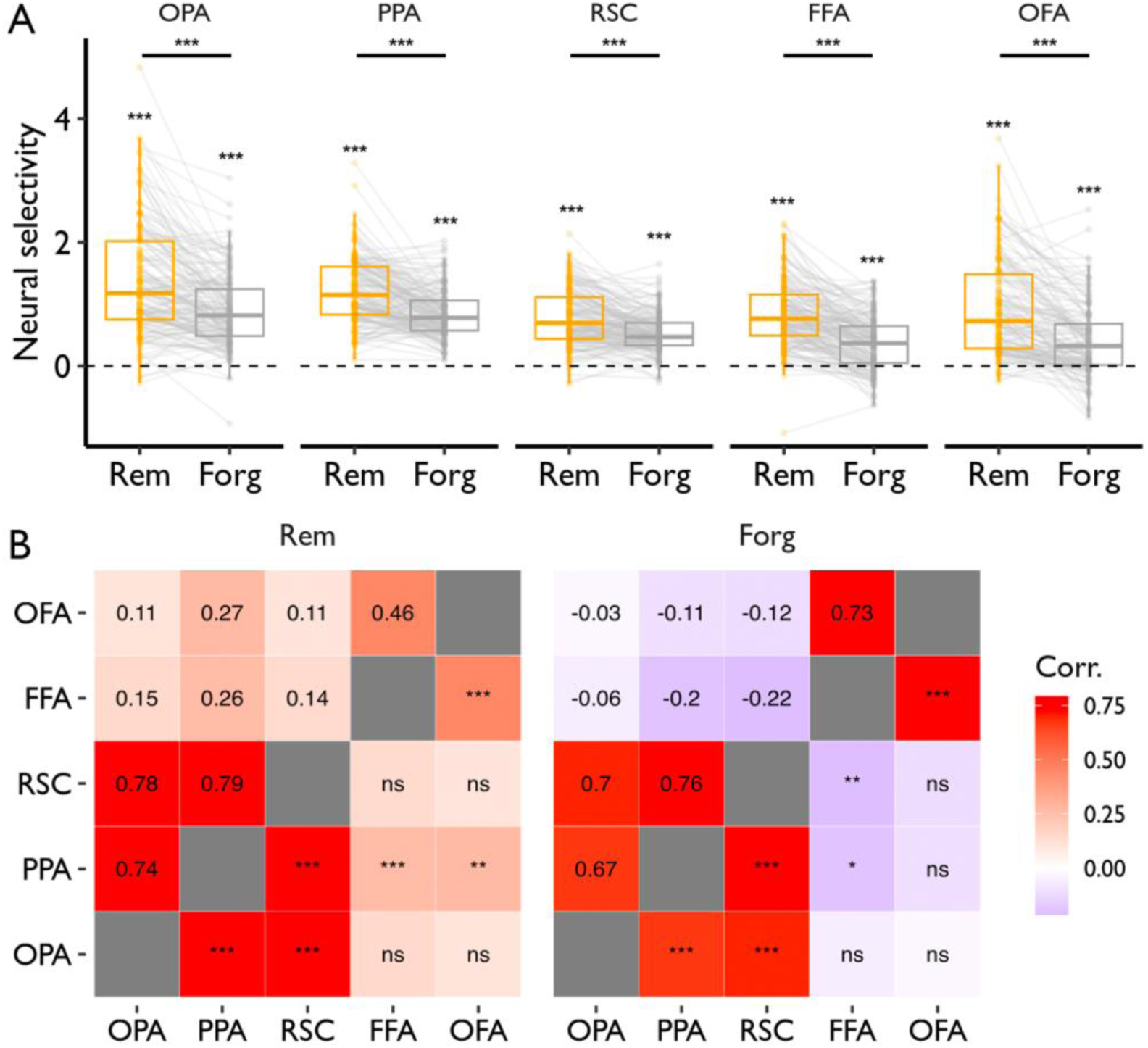
Neural selectivity varies with subsequent memory and remains more consistent within the same types of ROIs. (A) Neural selectivity in study-specific ROIs was greater for remembered trials than for forgotten trials. (B) Neural selectivity was correlated among place-(i.e., PPA, OPA, and RSC) or face-selective ROIs (i.e., FFA and OFA); the upper triangles show Pearson’s correlations, and the lower triangles show significance; ns = non-significant, *p < .05, **p < .01, ***p < .001.

**Fig. S4.**
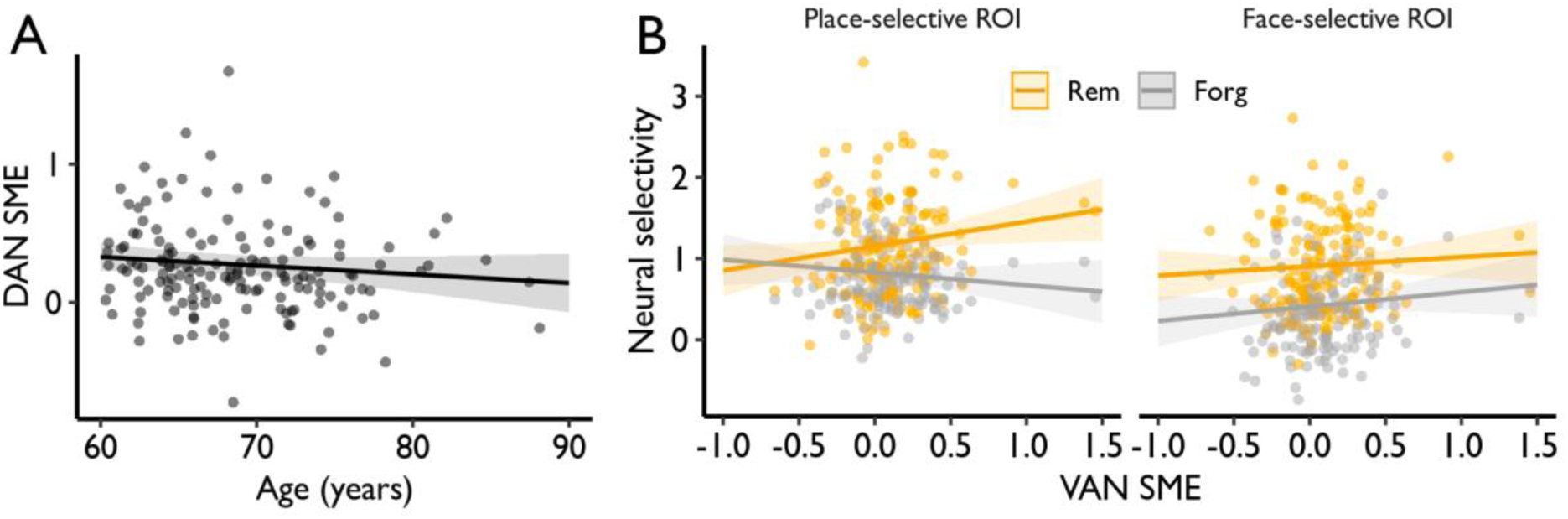
Relationship between DAN SME and age and neural selectivity. (A) Scatterplot of DAN SME by age. (B) Scatterplots of neural selectivity by VAN SME. DAN, dorsal attention network; VAN, ventral attention network; SME, subsequent memory effect.

**Fig. S5.**
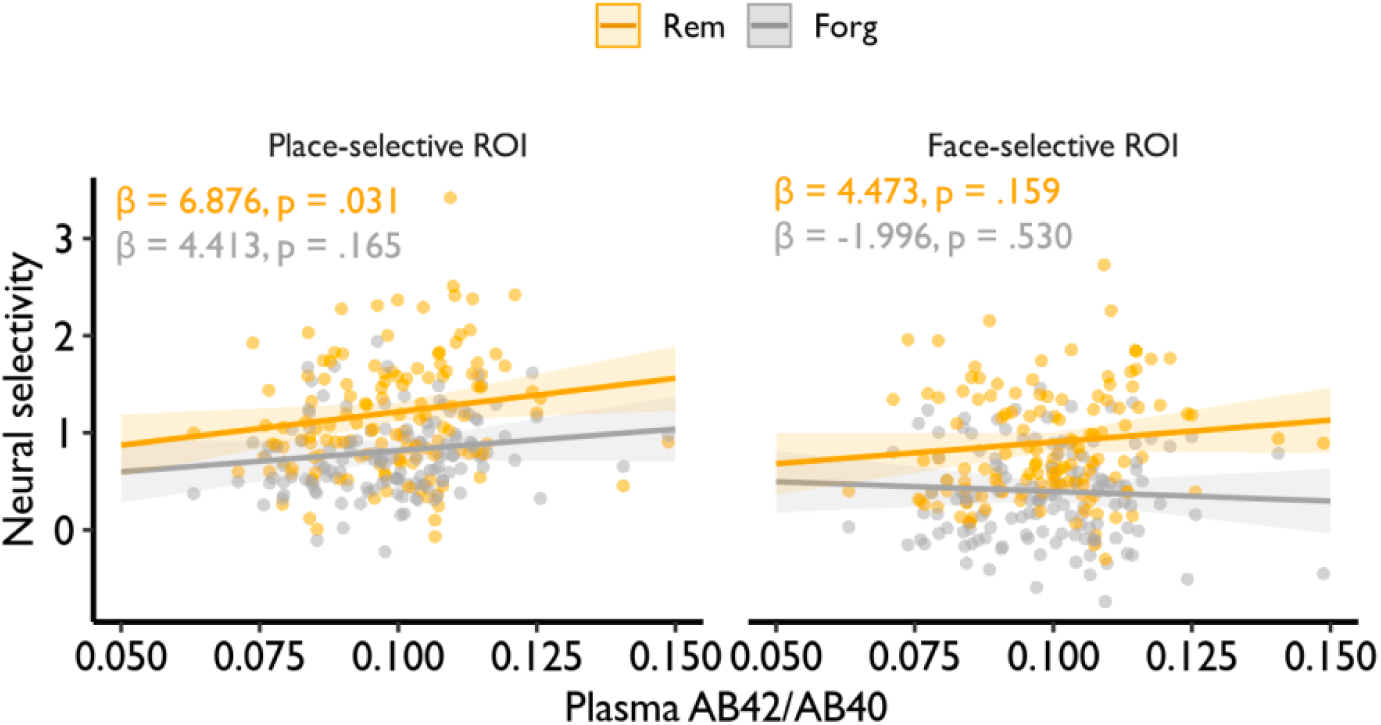
Relationship between neural selectivity and plasma Aβ_42_/Aβ_40_ (n = 136).

**Fig. S6.**
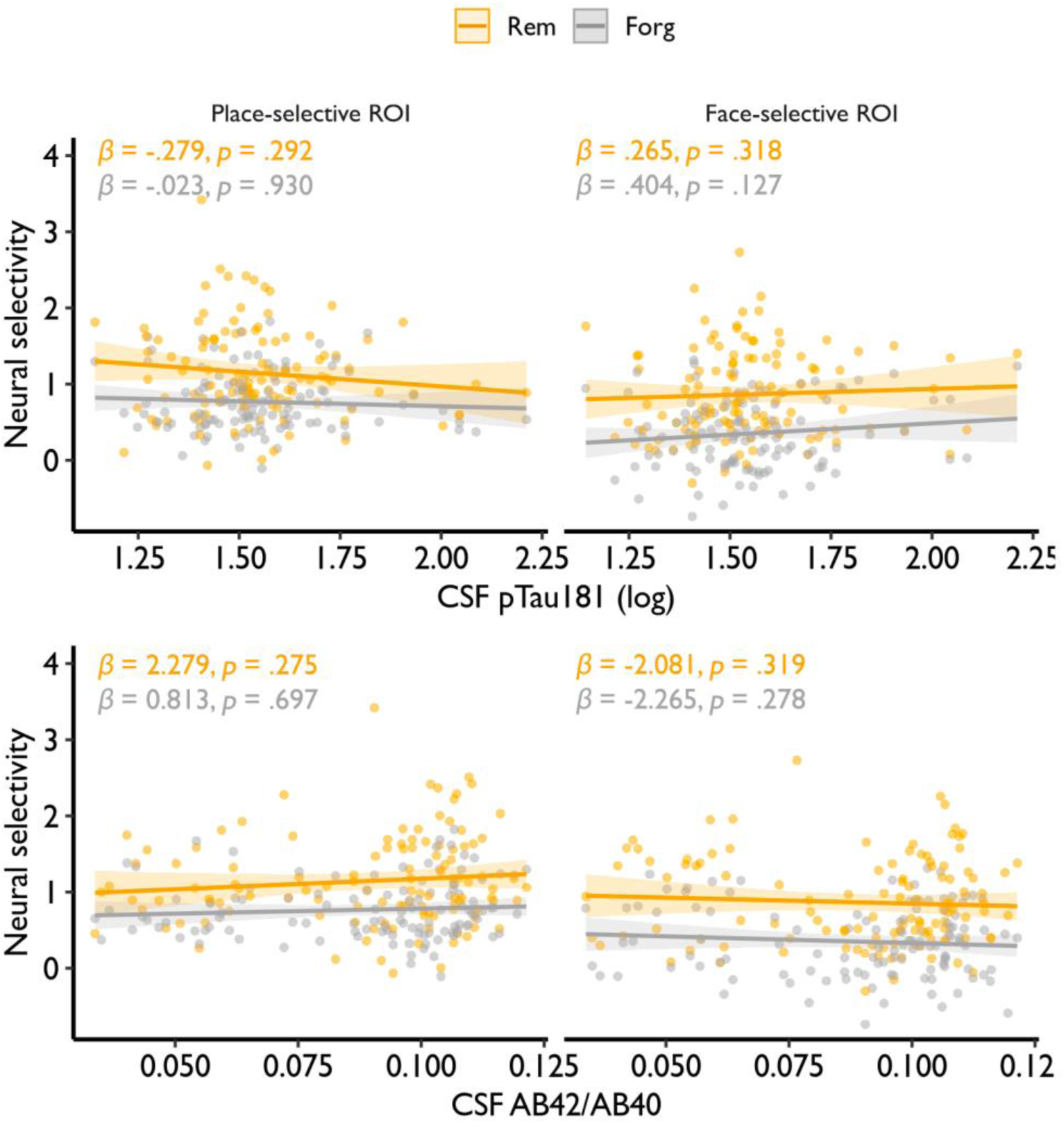
Relationship between neural selectivity and CSF AD biomarkers. (top) pTau_181_ and (bottom) Aβ_42_/Aβ_40_ (n = 115).

**Fig. S7.**
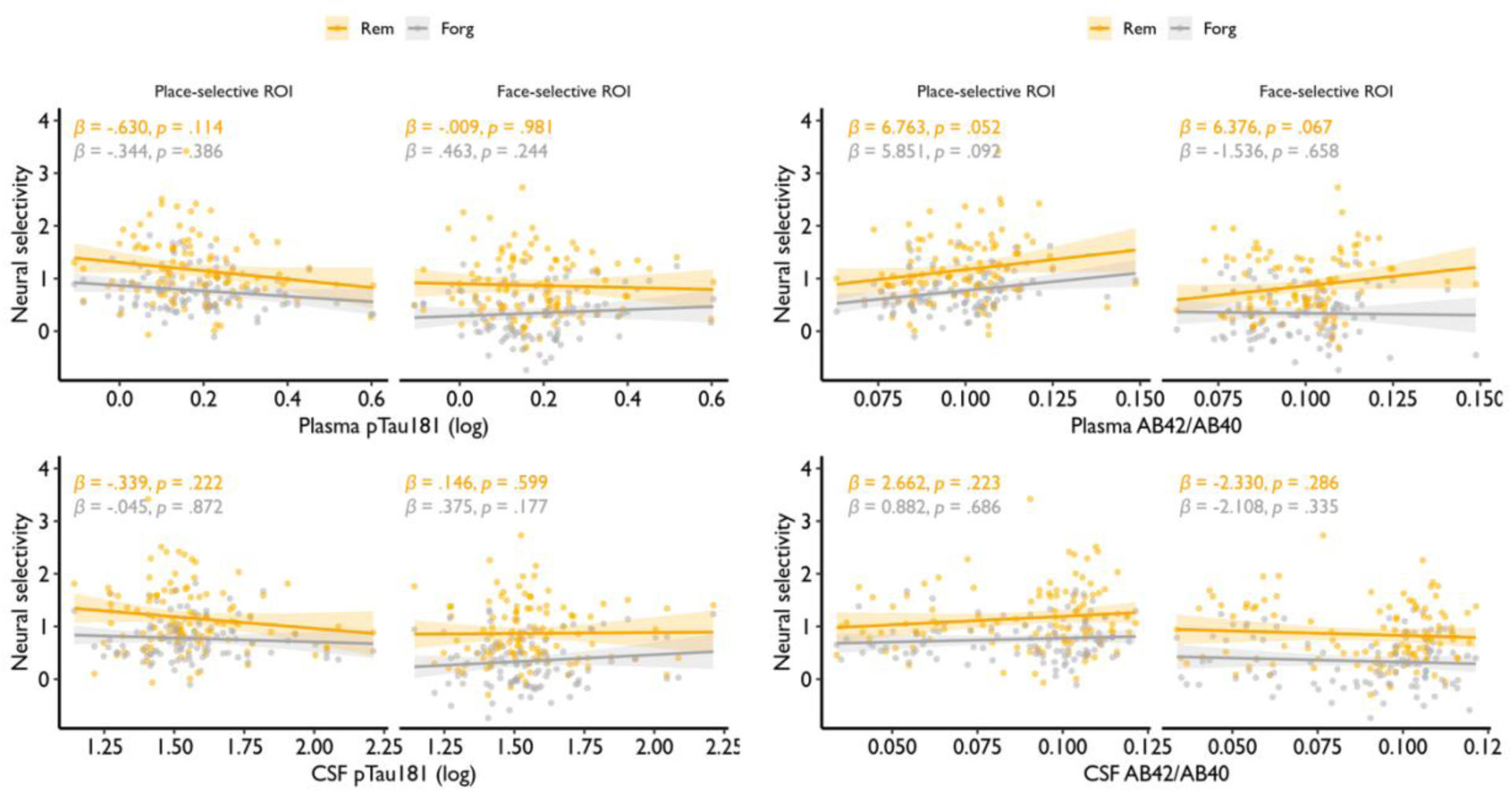
Relationship between neural selectivity and AD biomarkers in the subset of participants with both plasma and CSF. Neural selectivity is plotted as a function of pTau_181_ (n = 104) and Aβ_42_/Aβ_40_ (n = 103) in (top) plasma and (bottom) CSF.

**Fig. S8.**
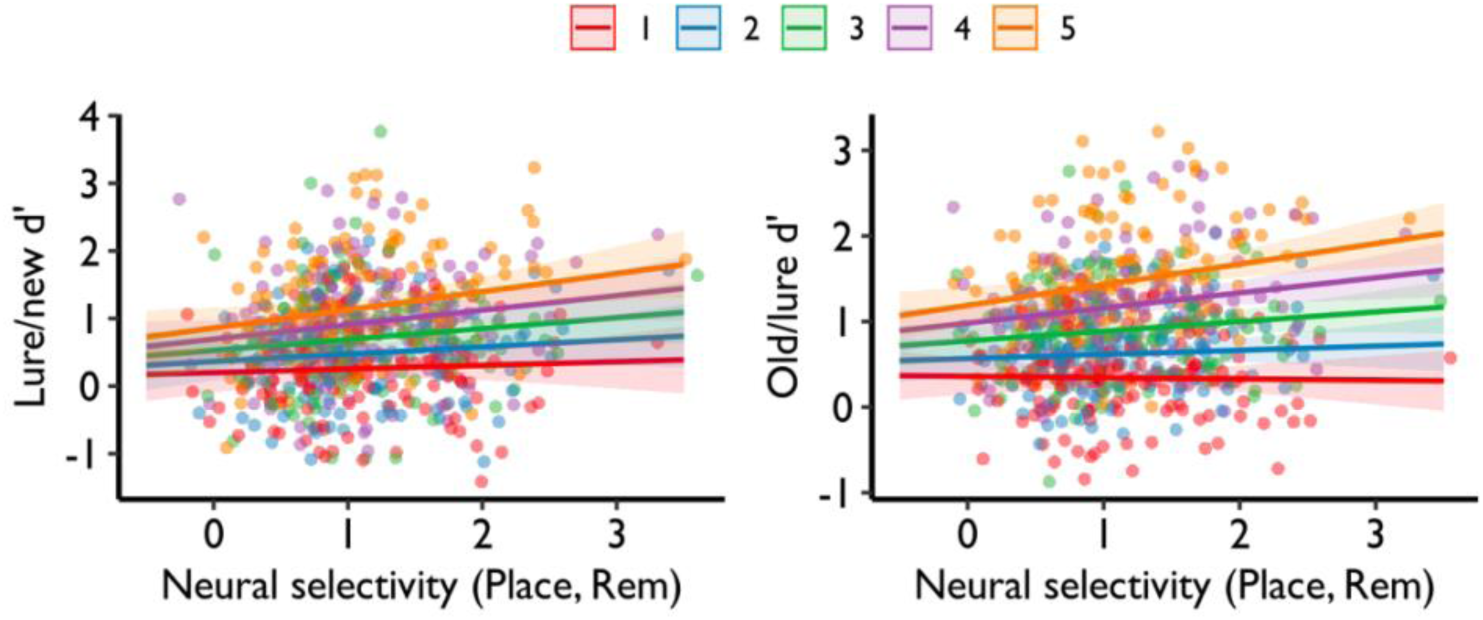
Neural selectivity interacts with target-lure similarity. MST performance varied as a function of neural selectivity and target-lure similarity, which ranged from high (1) to low (5) similarity.

**Fig. S9.**
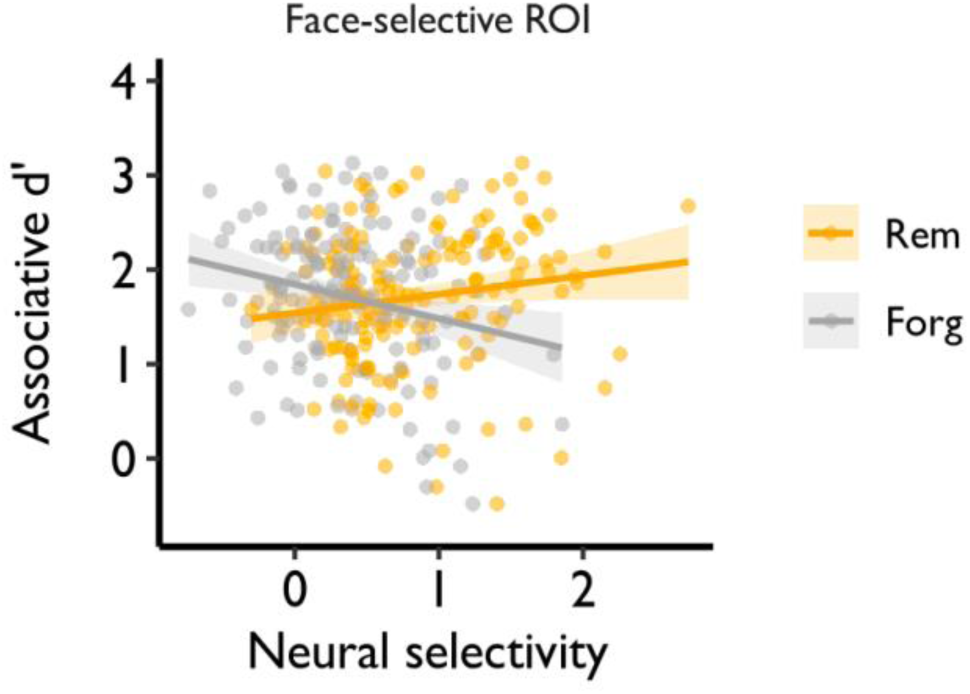
Relationship between neural selectivity in face-selective regions and overall associative *d’*.

